# Degeneracy in negative feedback (NFBL) and incoherent feedforward (IFFL) loops: Adaptation and resonance

**DOI:** 10.1101/2023.08.13.553122

**Authors:** Alejandra C. Ventura, Horacio G. Rotstein

## Abstract

Degeneracy in dynamic models refers to these situations where multiple combinations of parameter values produce identical patterns for the observable variable. We investigate this phenomenon in two qualitatively different adaptive circuit mechanisms: nonlinear feedback loop (NFBL) and incoherent feedback loop (IFFL). We use minimal models of these circuit types together with analytical calculations, regular perturbation analysis, dynamical systems tools and numerical simulations. In response to constant (or step-constant) inputs, NFBLs and IFFLs produce and overshoot allowing the observable variable to return to a value closer to baseline than the peak (adaptation). We identify the dynamic principles underlying the emergence of degeneracy in adaptive patterns both within and across circuit types in representative NFBL and IFFL models in terms of biologically plausible parameters. We identify the conditions under which degeneracy persists in response to oscillatory inputs with arbitrary frequencies, giving rise to resonance and phasonance degeneracy. This naturally extends to the response of adaptive systems to time-dependent inputs within a relatively large class. By using phase-plane analysis, we provide a mechanistic, dynamical systems-based interpretation of degeneracy. Our results have implication for the understanding of adaptive systems, for the relationship between adaptive and resonant/phasonant systems, for the understanding of complex biochemical circuits, for neuronal computation, and for the development of methods for circuit and dynamical systems reconstruction based on experimental or observational data.

## 1 Introduction

The study of adaptive responses to constant (or step-constant) inputs (Fig. 1-B, top) has been the focus of several studies in Systems Biology [1–6]. These patterns consists of an overshoot type of response, where the observable variable first increases and then decays towards a steady-state value (Fig. 1-C) [1]. We refer to these patterns as adaptive patterns and to the systems that produce them as adaptive systems.

**Figure 1:**
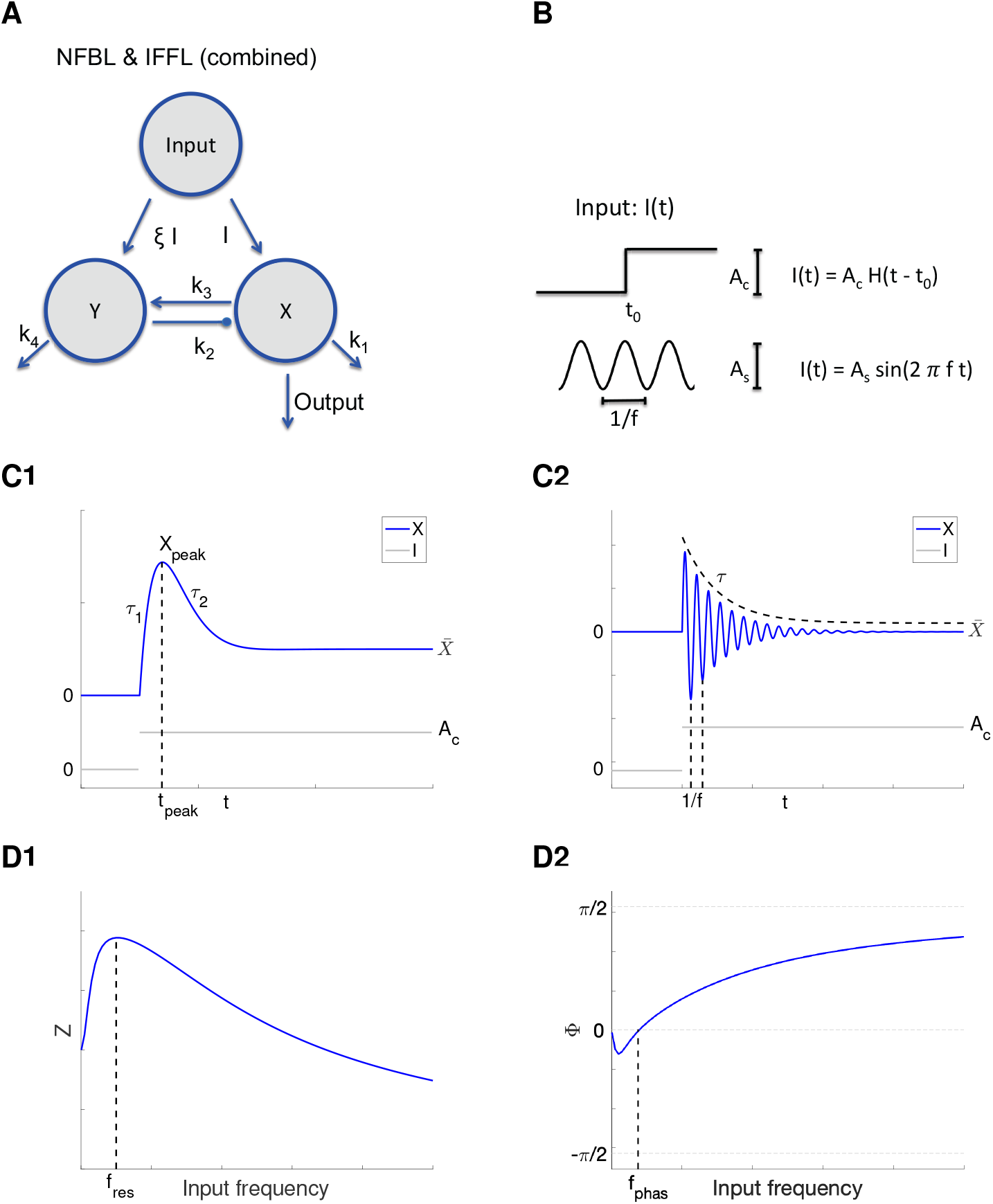
Negative feedback loop (NFBL) and incoherent feedforward loop (IFFL) schematic diagram. **A**. Combined NFBL and IFFL circuit. The circuit represents a NFBL for *ξ* = 0 and a IFFL for *k*_3_ = 0. The input arrives into the nodes *X* (with strength given by *I*) and *Y* (with strength given by *ξ I*). The node *X* activates the node *Y* (rate constant *k*_3_), which in turn inhibits the node *X* (rate constant *k*_2_). Both nodes exhibit self-regulation decay (linear decay rate constants *k*_1_ and *k*_4_). The output is measured in the node X. **B**. Representative input functions. Top: step-constant input function with amplitude *A*_*c*_. The jump occurs at *t* = *t*_0_. Bottom: sinusoidal input function with amplitude *A*_*s*_ and frequency *ω*. **C**. Representative response patterns of the observable variable *X* to a step-constant input *I* (light gray). **C1**. Representative adaptive pattern. In response to the abrupt change in *I* from *I* = 0 to *I* = *A*_*c*_, *X* first increases, reaches a peak *x*_*peak*_ at *t* = *t*_*peak*_, and then decreases towards a steady-state value 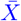. The three attributes of the response pattern are 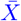 and the two time constants *τ*_1_ and *τ*_2_. **C2**. Representative damped oscillatory pattern. The three attributes of the response pattern are 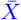, the frequency *ω* and the envelope time constant *τ*. **D**. Representative resonant (D1) and phasonant (D1) response profiles to oscillatory inputs. **D1**. Normalized amplitude response *Z* defined as the quotient of the steady state output amplitude and the input amplitude *A*_*s*_. The resonant frequency *f*_*res*_ is the input frequency at which *Z* peaks. **D2**. Phase (-shift) response Φ defined as difference between the peaks of the steady-state output amplitude and the input amplitude normalized by the period. The phasonant frequency *f*_*phas*_ is the (non-zero) input frequency at which Φ = 0.

**Figure 2:**
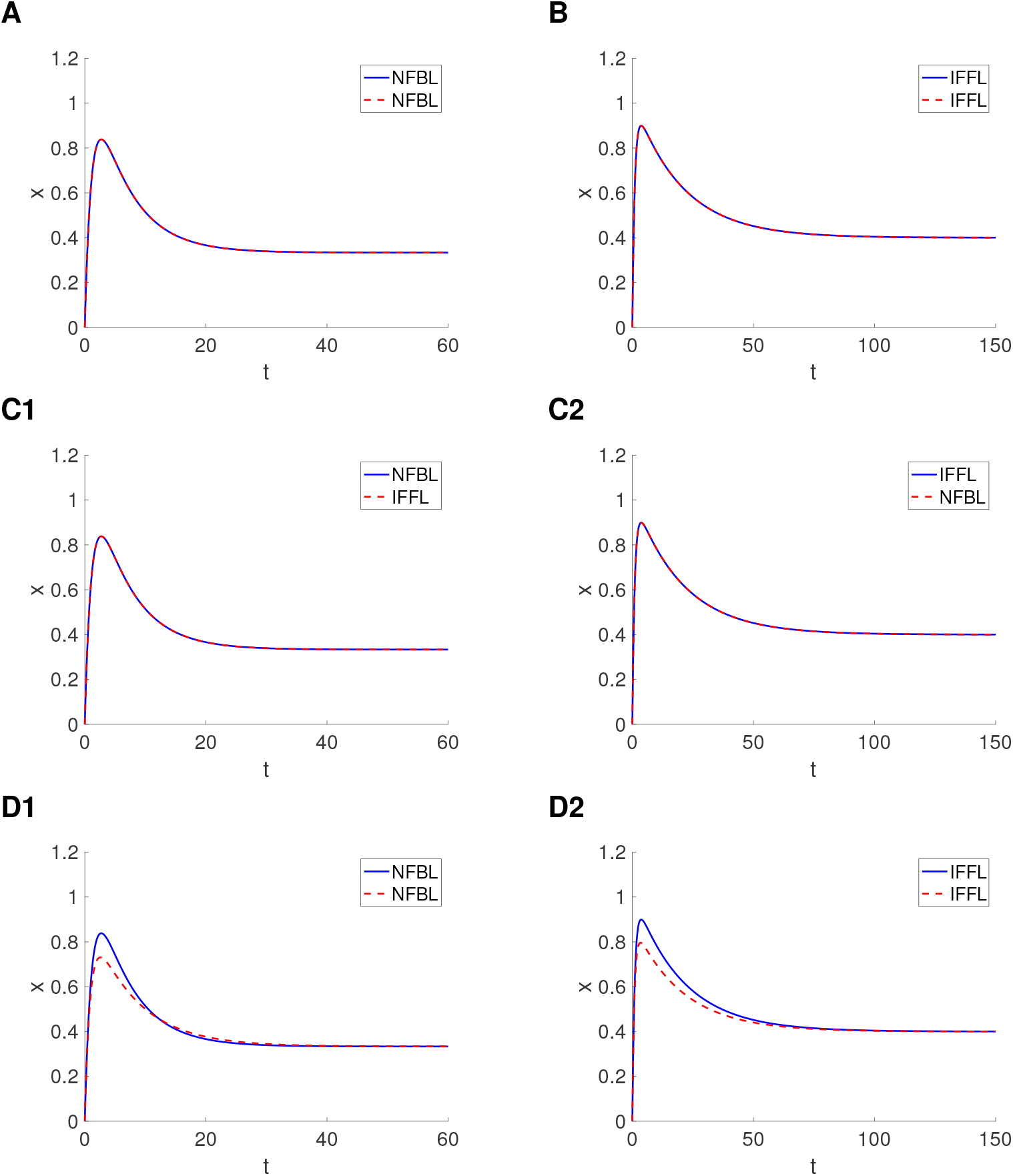
Degeneracy within and across circuit types: representative examples of adaptive responses (to a constant input). We used System (1)-(2) with *A*_*c*_ = 1, *A*_*s*_ = 0, *x*(0) = 0 and *y*(0) = 0. **A**. Degeneracy within NFBLs. Parameter values: *k*_1_ = 1, *k*_2_ = 2, *k*_3_ = 0.05, *k*_4_ = 0.05 (blue), *k*_1_ = 1, *k*_2_ = 1, *k*_3_ = 0.1, *k*_4_ = 0.05 (dashed-red). **B**. Degeneracy within NFBLs. Parameter values: *K*_1_ = 1, *K*_2_ = 2, *ξ* = 0.015, *K*_4_ = 0.05 (blue), *K*_1_ = 1, *K*_2_ = 1, *ξ* = 0.03, *K*_4_ = 0.05 (dashed-red). **C**. Degeneracy across NFBLs and IFFLs. Parameter values: *k*_1_ = 1, *k*_2_ = 2, *k*_3_ = 0.05, *k*_4_ = 0.05 (C1, blue), *K*_1_ = 0.8794, *K*_2_ = 1.2056, *ξ* = 0.1, *K*_4_ = 0.1706 (C1, dashed-red), *K*_1_ = 1, *K*_2_ = 2, *ξ* = 0.015, *K*_4_ = 0.05 (C2, blue), *k*_1_ = 1.03, *k*_2_ = 0.588, *k*_3_ = 0.05, *k*_4_ = 0.02 (C2, dashed-red). **D**. Balance, but not degeneracy within circuit types. Parameter values: *k*_1_ = 1, *k*_2_ = 2, *k*_3_ = 0.05, *k*_4_ = 0.05 (D1, blue), *k*_1_ = 1.2, *k*_2_ = 1.8, *k*_3_ = 0.05, *k*_4_ = 0.05 (D1, dashed-red), *K*_1_ = 1, *K*_2_ = 2, *ξ* = 0.015, *K*_4_ = 0.05 (D2, blue), *K*_1_ = 1.15, *K*_2_ = 1.8, *ξ* = 0.015, *K*_4_ = 0.05 (D2, dashed-red).

Biologically, the generation of adaptive patterns is determined by the interplay of the input and the system’s properties. From the dynamical systems point of view, the generation of adaptive patterns requires two-dimensional systems where the activity of an observable state variable is controlled or modulated by second, possibly hidden state variable. From the circuits point of view, the two variables can be embedded in a single node or in different nodes, depending on the biological context and conceptual framework.

Two minimal two-node circuit architectures have been identified as being able to produce adaptive behavior (Fig. 1-A) [3]: the negative feedback loop (NFBL, *ξ* = 0) and the incoherent feedforward loop (IFFL, *k*_3_ = 0). Adaptive patterns are the result of network regulation (processing) in NFBLs, while they are orchestrated by the input to the two nodes in IFFLs. Specifically, in the NFBL (*ξ* = 0), the observable node (X) receives the input (amplitude *A*_*c*_) and activates the inhibitor node (Y), which in turn inhibits X. In the IFFL (*k*_3_ = 0), in contrast, both the observable (X) and inhibitor (Y) nodes receive the inputs with an amplitude ratio equal to *ξ* and Y inhibits X, but Y is not activated X. These circuits have been associated to the processing of temporal information and related regulatory functions in a number of systems [7–12] (IFFL) and [13–19] (NFBL).

Dynamically, the generation of adaptive patterns in NFBLs and IFFLs requires that the observable evolves faster than the inhibitor so that the former can grow before being forced by the latter to turn around and decrease. These time scales, and the resulting properties of the adaptation peak, are determined by the combined effect of the model parameters and may differ for comparable NFBLs and IFFLs having identical parameter values except for these defining the type of circuit (*ξ* and *k*_3_). On the other hand, because balances between the activation and inhibition processes are expected, the question arises of whether NFBLs and IFFLs can produce identical adaptive patterns for multiple combinations of model parameters, thus giving rise to degeneracy [20, 21] both within and across circuit types. This in turn raises the question of whether and how the circuit properties can be inferred from the observable data and, in particular, under what conditions it is possible to distinguish between NFBLs and IFFLs from experimental or observational available data. Despite the pervasive presence of adaptive patterns in biological and related systems, these mechanistic issues are not well understood. In this paper we systematically address these questions in minimal models of NFBLs and IFFLs.

The dynamics of the nodes are assumed to be relatively simple: they are unidimensional and exhibit linear decay representing self-regulation (rates *k*_1_ and *k*_4_ in Fig. 1-A). We use linear and nonlinear models. Linear models are mathematically tractable and therefore allow us to gain intuition that can be used for the analysis of nonlinear models. The nonlinearities arise from the properties of the connectivity. We consider two types of nonlinearities: multiplicative, based on the law of mass action [22–24], and divisive (of sigmoidal type), based on the Michaelis-Menten formulation [25]. They are representative of the two ways inhibition can affect the rate of change of the observable variable. In the context of the diagram in Fig. 1-A, increasing values of the inhibitory variable (*y*) causes an increase of the negative contribution (negative term) to the rate of change of *x* in the multiplicative case and a decrease of the positive contribution (positive term) to the rate of change of *x* in the second case.

We analyze the nonlinear models by using dynamical systems tools (comparative phase-plane analysis). An important result of this analysis is that the mechanisms of generation of degeneracy in the linear and nonlinear comparable models (e.g., linear and nonlinear models of the same circuit types such as two NFBLs, two IFFLs, or one NFBL and one IFFL) are qualitatively similar.

Previous work by other authors [26] has focused on the discrimination between NFBLs and IFFLs, for example, by analyzing and comparing the circuits response to time-dependent signals such as periodic signals [4, 27] and ramps [28] or used stochastic models [29]. However, none of these approaches considered the presence of degeneracy in adaptive systems and the resulting adaptive patterns.

Common to these approaches is the use of time-dependent external signals, which can be captured by the circuits’ impedance amplitude and phase profiles in response to oscillatory inputs. Previous work has shown that adaptive systems can produce resonance (band-pass filters) (Fig. 1-D1) and phasonance (Fig. 1-D2) in the observable variable in response to oscillatory inputs [30–33] (see also [34–36]). Resonance is defined as the ability of a system to exhibit a peak in the amplitude response to oscillatory inputs at a preferred (resonant) non-zero input frequency. Phasonance is defined as the ability of a system to exhibit a zero-phase (phase-shift) response to oscillatory inputs at a preferred (phasonant) input frequency. Mechanistically, while resonance and phasonance have been observed in adaptive systems, it is not clear what aspects of adaptation control the generation of resonance and phasonance in NFBLs and IFFLs, and whether the generation of these patterns depend on the same balances that give rise to the corresponding adaptive patterns. The question arises of whether the degeneracy present in adaptive systems in inherited to their responses to oscillatory inputs. Whether and how resonant and phasonant patterns in NFBLs and IFFLs exhibit degeneracy has consequences for the ability of time-dependent inputs both deterministic and stochastic (e.g., Gaussian white noise) to be appropriate input signals to resolve degeneracy within and across NFBLs and IFFLs.

By including the relevant ingredients of NFBLs and IFFLs, the minimal models we use achieve a healthy balance between tractability and complexity that allows to identify the different types of degeneracy in response to constant and oscillatory inputs, identify the basic principles that govern the generation of degeneracy and related phenomena in NFBLs and IFFLs in response to both constant and time-dependent inputs, and understand the similarities and differences between the degenerate patterns observed in the two circuit types. Our results shed light on the mechanisms of generation of degeneracy in more complex NFBLs and IFFLs and have implications for the identification and development of the appropriate tools to differentiate between circuit types in the presence of degeneracy based on experimental or observational data. Additionally, the tools we use in this paper can be extended to the analysis of more complex models of NFBLs and IFFLs. We illustrate this by extending our work to include neuronal rate models describing the evolution of the instantaneous firing rates of excitatory and inhibitory networks [37–41].

## 2 Methods

### 2.1 Models of NFBL and IFFL circuits

#### 2.1.1 Linear NFBL and IFFL circuits

The linear NFBL and IFFL circuits (Fig. 1-A) are generically described by the following system

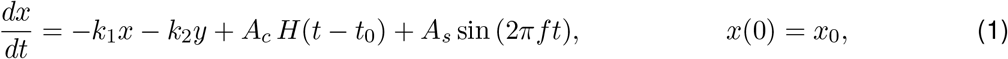

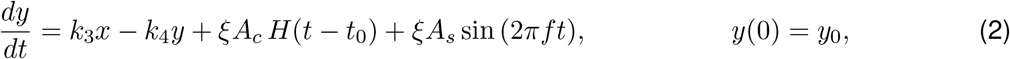

where *x* represents the observable, *y* represents the inhibitor, *k*_1_, *k*_2_, *k*_3_, *k*_4_, *A*_*c*_, *A*_*s*_, *f, ξ, x*_0_, *y*_0_ and *t*_0_ are non-negative constants, and *H*(*t*) is the Heaviside function. For the NFBL, *ξ* = 0, while for the IFFL, *k*_3_ = 0. The choice of parameters *x*_0_ = *y*_0_ = *t*_0_ = 0 and *A*_*c*_ *>* 0 corresponds to a system initially in equilibrium experiencing a sudden increase in the constant input at *t* = 0.

#### 2.1.2 Nonlinear NFBL and IFFL circuits

We consider NFBL and IFFL models with two types of nonlinearities in the inhibitory connectivity (from the node Y to the node X in Fig. 1-A): multiplicative and divisive. The general formulation is given by

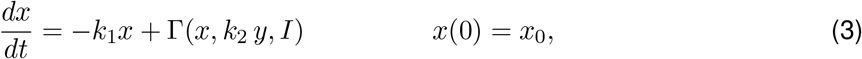

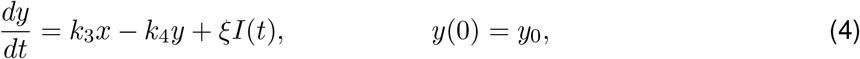

where

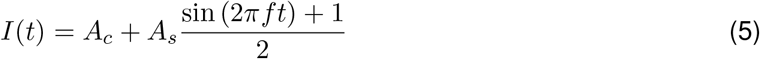

and all the parameters are as for the linear models in Section 2.1.1.

The multiplicative nonlinearity (M-) is based on the law of mass action [22–24]

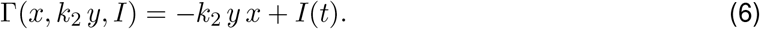

It produces a multiplicative interaction between the observable and the inhibitor. Increasing values of *y* increase the negative contribution of the nonlinear term to the rate of change of *x*.

The divisive nonlinearity is based on the Michaelis-Menten formulation [25] (see also [42] for covalent modification cycles). We consider two different forms, which differ on the location of the input signal

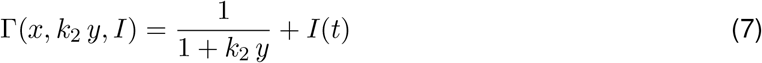

and

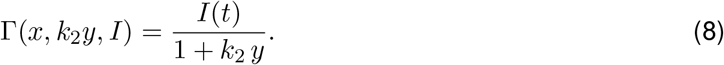

In both cases, increasing levels of values of *y* decrease the positive contribution of the nonlinear term to the rate of change of *x*. We refer to these two forms as the divisive nonlinearity with an additive (Da-) and multiplicative (Dp-) inputs, respectively. We refer to the NFBL and IFFL models having these three types of nonlinearities as the M-, Da- and Dp-NFBL and -IFFL models.

### 2.2 Model attributes and observable attributes: The linear case

By observable attributes we refer to the parameters describing the circuit’s response to a given input or model solution (e.g., Fig. 1-C). For a linear system, at least one set of observable attributes can be analytically calculated from the solution, while for nonlinear systems they need to be numerically computed from the response patterns. By model attributes we refer to the reduced parameters, consisting of a combination of the (original) model parameters obtained when simplifying the model. For linear systems, the observable attributes can be computed from the model attributes and the number of observable and model attributes coincide.

In order to introduce these ideas, we use the following generic linear system that generalizes the NFBL and IFFL linear models described above

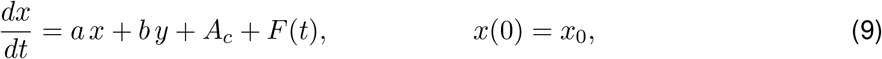

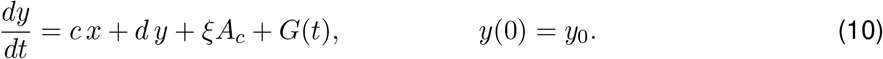

The model parameters *a, b, c, d, ξ* and *A*_*c*_ are assumed to be constant. For this system to represent the NFBL or the IFFL, *a* = −*k*_1_ *<* 0, *b* = − *k*_2_ *<* 0, *c* = *k*_3_ ≥0, *d* = −*k*4 *<* 0 and *ξ*≥ 0. Additionally, for the NFBL, *ξ* = 0, while for the IFFL, *c* = 0.

#### 2.2.1 Reduction to a second order linear ODE: Identification of the model attributes

System (9)-(10) can be rewritten as a linear second order ODE in terms of the variable *x* [43]

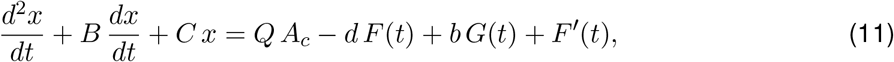

where the model attributes are given by

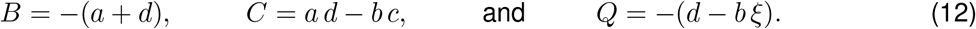

We refer to the reduced parameters *A, B* and *Q* in (12) as model attributes to differentiate them from the model parameters in System (9)-(10). In the absence of time-dependent inputs (*F* = *G* = 0), the number of parameters that effectively govern the dynamics of *x* is reduced from five (model parameters) to three (model attributes).

We use this model reduction to identify the degeneracy present in the NFBL and IFFL systems. As we show in this paper, other types of model reductions such the widely used non-dimensionalization [44–47] are too stringent to capture the degeneracy present in the NFBL and IFFL systems.

An alternative reduction consists of rewriting System (9)-(10) as a linear second order ODE in terms of the variable *y*

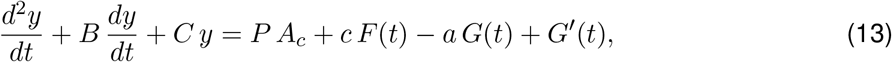

where the model attributes are given by *A* and *B* in (12) and

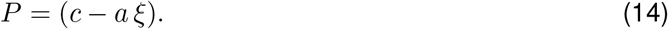

Note. The initial conditions are *x*(0) = *x*_0_ and *dx/dt*(0) = *a x*_0_ + *b y*_0_ + *A* + *F* (0) in eq. (11) and *y*(0) = *y*_0_ and *dy/dt*(0) = *c x*_0_ + *d y*_0_ + *ξ A* + *G*(0) in eq. (13).

#### 2.2.2 Computation of a set of observable attributes

Solutions to second order ODEs of the form

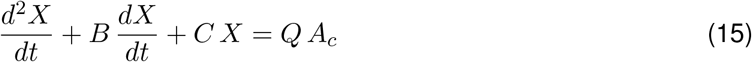

where *B, C, A*_*c*_ are constant are characterized by three observable attributes (in addition to the initial conditions) that can be extracted from the response patterns to the constant input *A*_*c*_ (Fig. 1-C) following standard procedures to solve linear differential equations [43].

One observable attribute is the steady-state

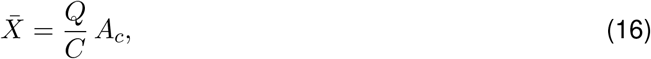

which is assumed to be stable. If the solution evolves monotonically or exhibits and overshoot ^1^, the other two observable attributes are the time constants

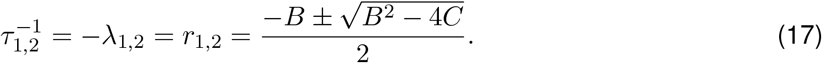

In this case, the eigenvalues *r*_1,2_ are real and negative. (Overshoots are often refer to as sags if the solution first decreases and then increases.). If, instead, the solution exhibits damped oscillations ^2^ (*r*_1,2_ are complex with non-zero imaginary part), the two attributes are the frequency and the oscillations envelope time constant

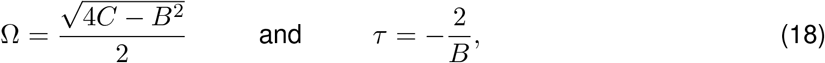

respectively. We refer to them as observable attributes to differentiate them from the model attributes (*B, C* and *Q*).

The model attributes (*B, C* and *Q*) can be uniquely computed from the observable attributes (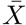, *τ*_1,2_, or, alternatively, 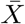, *τ* and *ω*) (see Supplementary Material).

We note that these are not the only possible set of attributes that can be extracted from the response patterns (e.g., the time constants *τ*_1,2_ can be substituted by the peak and peak time if the solution exhibits an overshoot), but the total number of attributes that completely determine the solution *X* is three. We also note that when the time constants are not different enough (the time scales are not well separated) two attributes are enough to approximately describe the solution.

### 2.3 Response of two-dimensional systems to oscillatory inputs

The response of a two-dimensional system to oscillatory inputs can be characterized by the impedance amplitude (*Z*) and phase (Φ) profiles. These are curves of the impedance amplitude (illustrated in Fig. 1-D1) and phase-shift (illustrated in Fig. 1-D2) as a function of the input frequency. For linear systems, the input and output frequencies coincide. For the nonlinear models we consider in this paper, the number of input and output cycles coincide and the response amplitude is uniform across cycles for a given input frequency (assuming steady-state).

The quantities *Z* and Φ are the real and imaginary parts of the complex impedance **Z**, defined as the quotient of the Fourier transforms of the output and input signals. For linear systems, *Z* and Φ can be computed analytically. For the nonlinear models we use in this paper, we use a generalization of the impedance for linear system [33].

The notion of impedance is borrowed from the field of electric circuits. While the interpretation of the responses of biological and electric systems to oscillatory signals is inherently different, we use the same terminology for convenience.

#### 2.3.1 Linear systems: Impedance amplitude and phase profiles

We consider the two-dimensional linear system described by eqs. (9)-(10) in Section 2.2 with

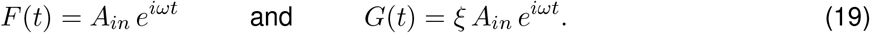

The stationary solution of system (9)-(10) is given by

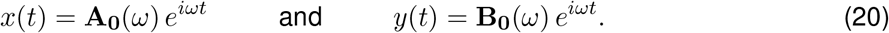

By analogy with electric circuits, we define the (complex) impedances

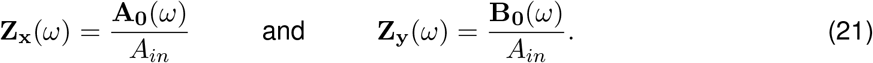

We present the detailed calculations in the Supplementary material.The impedance amplitudes are given by

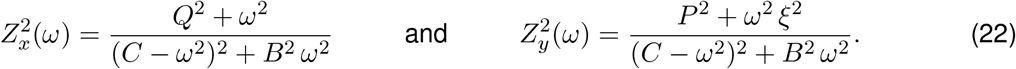

The impedance phases (phase-shift) are given by

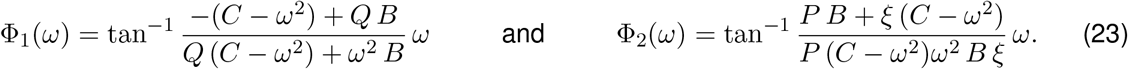

#### 2.3.2 Extension of the notions of impedance amplitude and phase profiles to nonlinear systems

We extend the notion of *Z* and Φ to characterize the response of the observable variable *X* in a nonlinear system receiving an oscillatory input of the form (5). Following [33], we define

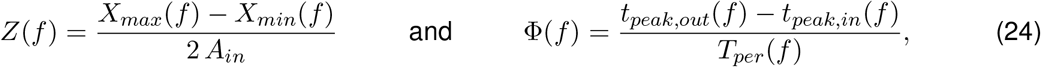

where *X*_*max*_(*f*) and *X*_*min*_(*f*) are the maximum and minimum of the (stationary) oscillatory response, respectively, *t*_*peak,in*_(*f*) and *t*_*peak,out*_(*f*) are the peak times of the input and output signals, respectively, and *T*_*per*_(*f*) is the oscillation period.

The two quantities in (24) generalize the notion of impedance for linear systems described above under the assumption that the number of input and output cycles per unit of time coincides and the output voltage waveforms for each each input frequency are identical across cycles (assuming steady state). This is the case for linear models and a number of nonlinear models [32, 33, 49, 50]. We note that the nonlinearities are expected to affect the response waveforms as compared to the analogous responses of linear systems and the waveforms may significantly differ across frequencies.

## 3 Results

### 3.1 Degeneracy within and across circuit types in linear NFBLs and IFFLs in response to constant inputs

System (1)-(2) with *A*_*s*_ = 0 and *t*_0_ = 0 (without loss of generality) can be rewritten as a linear second order ODE of the form (15) in terms of the variable *x* [43] (see Section 2.2.1) where the model attributes are given by

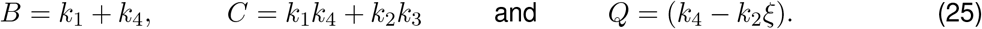

The response is completely determined by the the three observable attributes: the steady-state 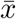 (16) and the time constants *τ*_1_ and *τ*_2_ (17) (e.g, Fig. 1-C1) or the oscillations frequency and envelope time constant (18) (e.g, Fig. 1-C2), which are uniquely determined by the model attributes.

Response patters having the same model or observable attributes for the same value of *A*_*c*_ are identical. Therefore, if one only has access to the observable variable *x*, as is often the case, the solutions are degenerate.

For the NFBL (*ξ* = 0),

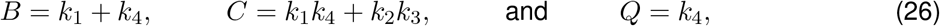

where as for the IFFL (*k*_3_ = 0),

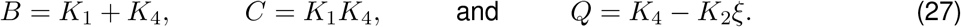

We use capital letters here for the rate constants in the IFFL to indicate that the model parameters in the NFBL and IFFL models for which the model attributes (*B, C* and *Q*) are the same, have not necessarily the same values.

#### 3.1.1 Degeneracy within circuit types

For the NFBL, from (26), *k*_1_ and *k*_4_ are uniquely determined by *B* and *Q*. The intra-circuit (NFBL) degeneracy is given by the multiple combinations of *k*_2_ and *k*_3_ that can produce

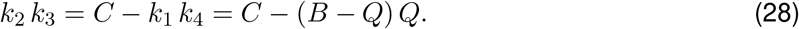

We illustrate this in Fig. 7-A.

For the IFFL, from (27), *K*_1_ and *K*_4_ are uniquely determined by *B* and *C*. The intra-circuit (IFFL) degeneracy is given by the multiple combinations of *K*_2_ and *ξ* that can produce

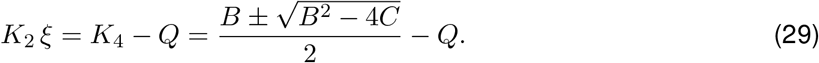

We illustrate this in Fig. 7-B.

Degeneracy in both circuit types reflects the multiple possible balances between the activation of *y* (*k*_3_ or *ξ*) and the inhibition *y* exerts on *x* (*k*_2_). There are other possible balances in both circuit types, but they will affect the shape of the response and therefore will not result in degeneracy (Fig. 7-D). For example, one can find multiple combinations of parameter values, other than exclusively the degenerate ones mentioned above, that keep 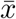 constant. Since this will necessarily require different values of *B*, then it will also require different values of the other two observable attributes.

#### 3.1.2 Degeneracy across circuit types

The cross-circuit degeneracy between the NFBLs and IFFLs follows from our discussion above. In practice, we first fix the parameters of the NFBL model and determine the values of *B, C* and *Q* from eqs. (26). Then, we find the values of the parameters of the IFFL model from eqs. (27) and (29). Alternatively, we fix the parameters of the IFFL model and the values of *B, C* and *Q* from eqs. (27). Then, we find the values of the parameters of the IFFL model from eqs. (26) and (28). We illustrate this in Fig. 7-C.

#### 3.1.3 Degeneracy in the observable variable (*x*) does not imply degeneracy in the hidden variable (*y*)

System (1)-(2) with *A*_*s*_ = 0 and *t*_0_ = 0 (without loss of generality) can be rewritten as a linear second order ODE of the form (13) (*F* = *G* = 0) in terms of the variable *y* [43] where *B* and *C* are given by (25) and

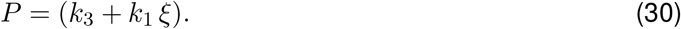

An analysis similar to the one carried above identifies the degeneracy structure for the hidden variable *y* and shows that this structure is different from the degeneracy structure for the variable *x*. This is because the right hand side term in eq. (13) (*F* = *G* = 0) involves the model attribute *P* instead of the model attribute *Q*.

### 3.2 The intra- and cross-circuit-type degeneracy in linear NFBLs and IFFLs persists in response to oscillatory inputs

The presence of degeneracy in biological circuits presents an obstacle in identifying the circuit architecture from observable data and distinguishing between NFBLs and IFFLs. Different types of time-dependent inputs have been used in the past to achieve the latter goal [4, 26–29]. In order to test the extent to which time-dependent inputs are appropriate tools to resolve the degeneracy within and across circuit types, here we examine the response of the NFBL and IFFL circuits to oscillatory inputs. We use eqs. (1)-(2) with *A*_*s*_ = 0 (and *A*_*c*_ = 0 without loss of generality).

The steady-state response for the observable variable is given by

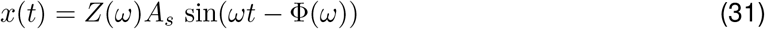

where *ω* = 2*πf* and the impedance amplitude and phase (see Section 2.3.1) are given, respectively, by eqs. (22) and (23).

Both quantities involve only the model attributes *B, C* and *Q* given by eqs. (25). It follows that the degeneracy structure of the response to oscillatory signals is identical to the degeneracy structure discussed in Section 3.1 for the response to constant inputs. Fig. 3 shows representative examples of the impedance amplitude and phase profiles (curves of the impedance amplitude and phase as a function of the input frequency) for the same parameter values as in Fig. 7-C. Therefore, oscillatory signals and, more generally, a large class of time-dependent signals are not useful tools to resolve the degeneracy for the linear adaptive systems discussed above.

**Figure 3:**
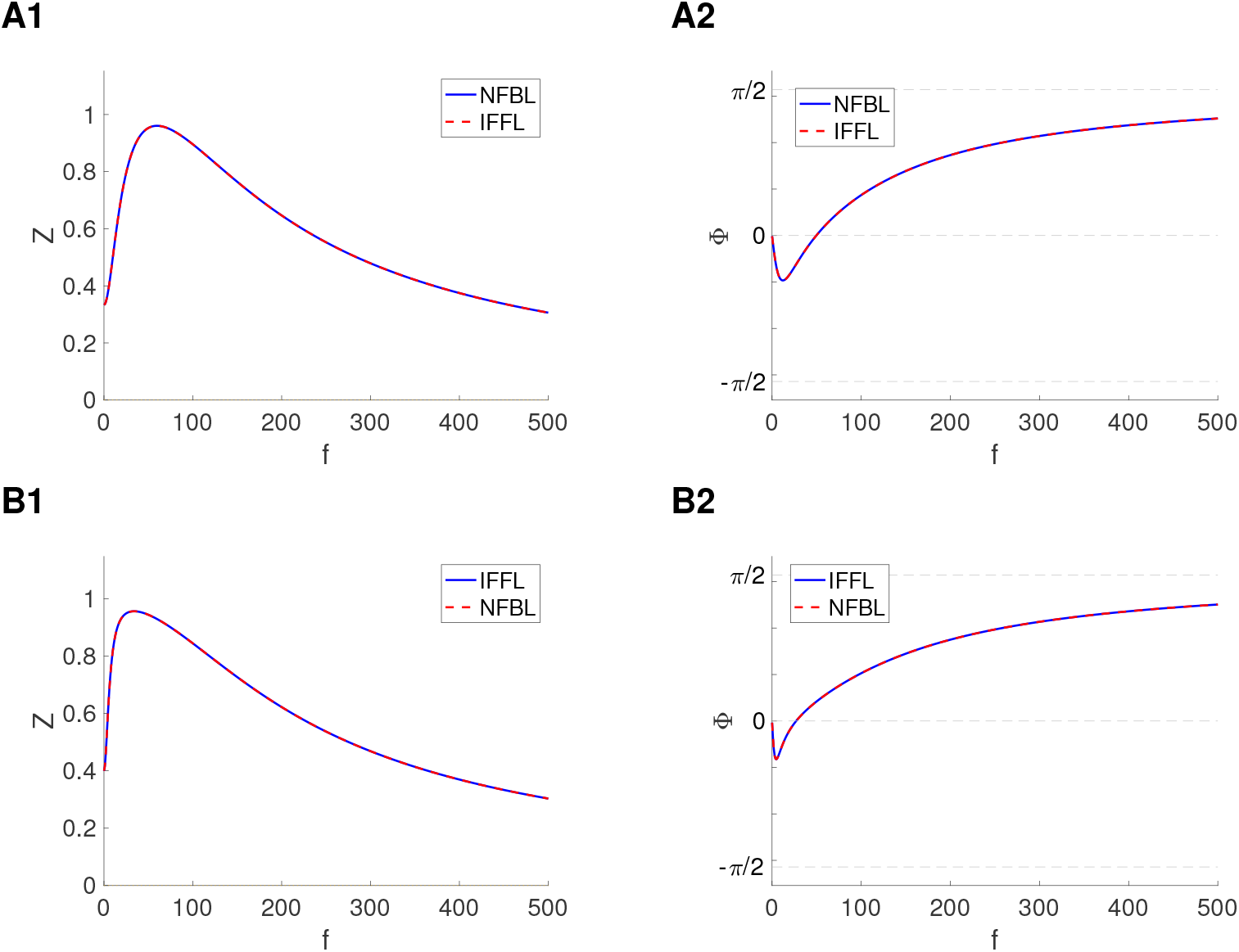
Degeneracy across circuit types: representative examples or resonant and phasonant profiles (in response to oscillatory inputs). We used System (1)-(2) with *A*_*c*_ = 0, *A*_*s*_ = 1, *x*(0) = 0 and *y*(0) = 0. **Left column**. Impedance amplitude profile. **Right column**. Phase profile. We use the same parameter values as in Fig. 7-C. **A**. We fixed the parameter values in the NFBL and computed the degenerate parameter values in the IFFL. Parameter values: *k*_1_ = 1, *k*_2_ = 2, *k*_3_ = 0.05, *k*_4_ = 0.05 (blue), *K*_1_ = 0.8794, *K*_2_ = 1.2056, *ξ* = 0.1, *K*_4_ = 0.1706 (dashed-red). **B**. We fixed the parameter values in the IFFL and computed the degenerate parameter values in the NBFL. Parameter values: *K*_1_ = 1, *K*_2_ = 2, *ξ* = 0.015, *K*_4_ = 0.05 (blue), *k*_1_ = 1.03, *k*_2_ = 0.588, *k*_3_ = 0.05, *k*_4_ = 0.02 (dashed-red).

### 3.3 Degeneracy within and across circuit-types is not captured by non-dimensionalized linear NFBLs and IFFLs

In the previous sections we identified the degeneracy present within and across NFBLs and IFFLs by using the model reduction technique described in Section 2.2.1. Another, widely used model reduction technique is non-dimensionalization [44–47]. By defining dimensionless variables and parameters (in terms of the original, dimensional ones) the complexity of models is reduced (the number of dimensionless, free parameters is smaller than the number of dimensional parameters), the magnitude of the free parameters is revealed, and the structure (e.g., temporal) of the solutions is easier to uncover than in the original model. The solutions to the (original) dimensional model can be obtained from the solutions to the non-dimensional model by reintroducing the (dimensional) model variables and parameters, and therefore non-dimensionalization does not result in any loss of information about the dynamics of the original model. For these reason, modelers often restrict their investigation to the simpler dimensionless models.

While non-dimensionalization reveals important structural properties of NFBLs and IFFLs as well as structural differences between them, the question arises of whether it captures the degeneracy present in these models both within and across circuit types.

#### 3.3.1 Non-dimensionalization of the NFBL and IFFL models

In order to address this, we define the following dimensionless variables

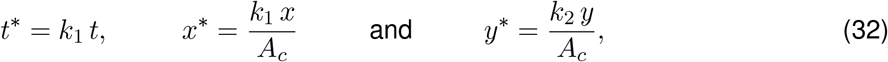

and parameters

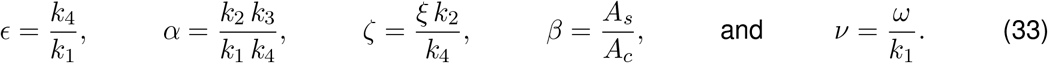

Substitution into system (1)-(2) yields

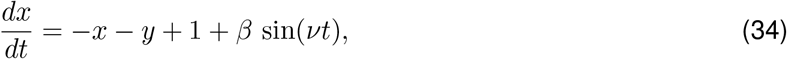

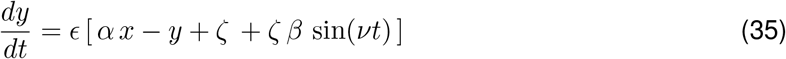

where we have dropped the “^*^” sign from the variables.

As before, system (34)-(35) with *β* = 0 can be rewritten as a linear second order ODE of the form

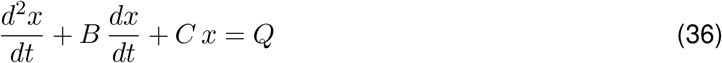

where

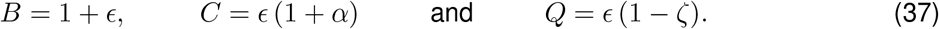

For the NFBL (*ζ* = 0),

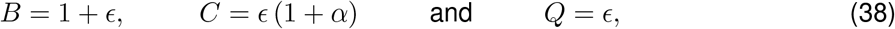

while for the IFFL (*α* = 0),

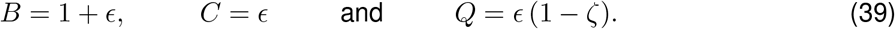

#### 3.3.2 The non-dimensional model highlights fundamental properties of NFBLs and IFFLs and differences between them

Some important differences between the NFBL and IFFL are immediately apparent from the non-dimensional model.

##### IFFL, but not NFBL can produce perfect adaptation

Perfect adaptation refers to the scenarios where the response pattern to a step-constant input returns to its baseline value prior to the arrival of the signal (*x* = 0 for the representative cases we are considering here). The steady-state solution to system (34)-(35) (*β* = 0) is given by 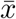 by eq. (16) with *A*_*c*_ = 1

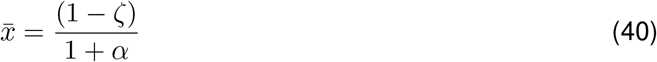

For the IFFL (*α* = 0), perfect adaptation is achieved when *ζ* = 1. For the NFBL (*ζ* = 0), achieving perfect adaptation requires *α* → ∞.

For the IFFL, perfect adaptation is possible because of the orchestration of activation and inhibition to the node X by the input signal. For the NFBL, the inhibitory node Y is not directly activated, but is activated by the node X. We note that for the linear IFFL 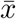 can become negative (*ζ >* 1), posing a limitation to the validity of the model.

##### NFBL, but not IFFL can produce intrinsic oscillations

The time constants *τ*_1_ and *τ*_2_ are given by (17)

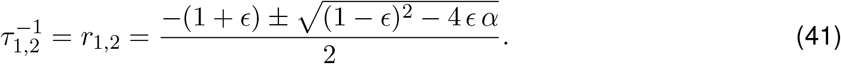

Adaptive solutions require the discriminant to be positive. For the NFBL (*ζ* = 0), the discriminant can become negative and therefore, intrinsic oscillations, with frequency and envelope time constant given by eq. (18), are possible. In contrast, for the IFFL (*α* = 0, the discriminant is always positive and therefore intrinsic oscillations are not possible. Intuitively, the IFFL is lacking a negative feedback.

##### IFFLs and NFBLs can exhibit resonance in the absence of intrinsic oscillations, but they do so by different mechanisms

It is often believed that resonance is an amplification of the response of a system at or around the natural frequency of the system receiving the oscillatory inputs. While resonance in physical systems (e.g., mechanical, electrical) is typically study in harmonic and damped oscillators [43], non-oscillatory systems can also exhibit resonance in the presence of negative feedback effects [30, 49]. This is the case of the NBFLs and IFFLs discussed in Section 3.2. NFBLs fall into the category of models investigated in [30,49]. In contrast, IFFLs do not have a negative feedback loop. Resonance results from a different mechanism where the preferred frequency response is the result of an oscillatory balance produced by the same type of orchestration of activation and inhibition to the node X described above for adaptation.

#### 3.3.3 Intra-circuit degeneracy is absent in the non-dimensionalized linear NFBLs and IFFLs

From (38) and (39), intra-circuit-type degeneracy is absent in both the NFBL and IFFL dimensionless models since the number of dimensionless parameters in each circuit type is smaller than the number of attributes. The non-dimensional models lack the necessary flexibility to capture the degeneracy present in the dimensional models since the non-dimensional model attributes are constrained: *B* = 1 + *Q* for the NFBL and *B* = 1 + *C* for the IFFL.

#### 3.3.4 Cross-circuit degeneracy is absent in the non-dimensionalized linear NFBLs and IFFLs

By comparing (38) and (39), the cross-circuit-type degeneracy is also absent in the dimensionless models. A fix value of the model attribute *B* determines *ϵ* in both the NFBL and IFFL. However, the equalities of *C* and *Q* between the NFBL and IFFL cannot be met since *α >* 0 (for C in the NFBL) and *ζ >* .0 (for *Q* in the IFFL). Multiple combinations of *α* and *ξ* can produce the same stationary value 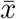. However, the time constants *τ*_1,2_ are different in the two models for *α >* 0 (Fig. 4-A). These differences are translated to the impedance and phase profiles (Fig. 4-B and -C).

**Figure 4:**
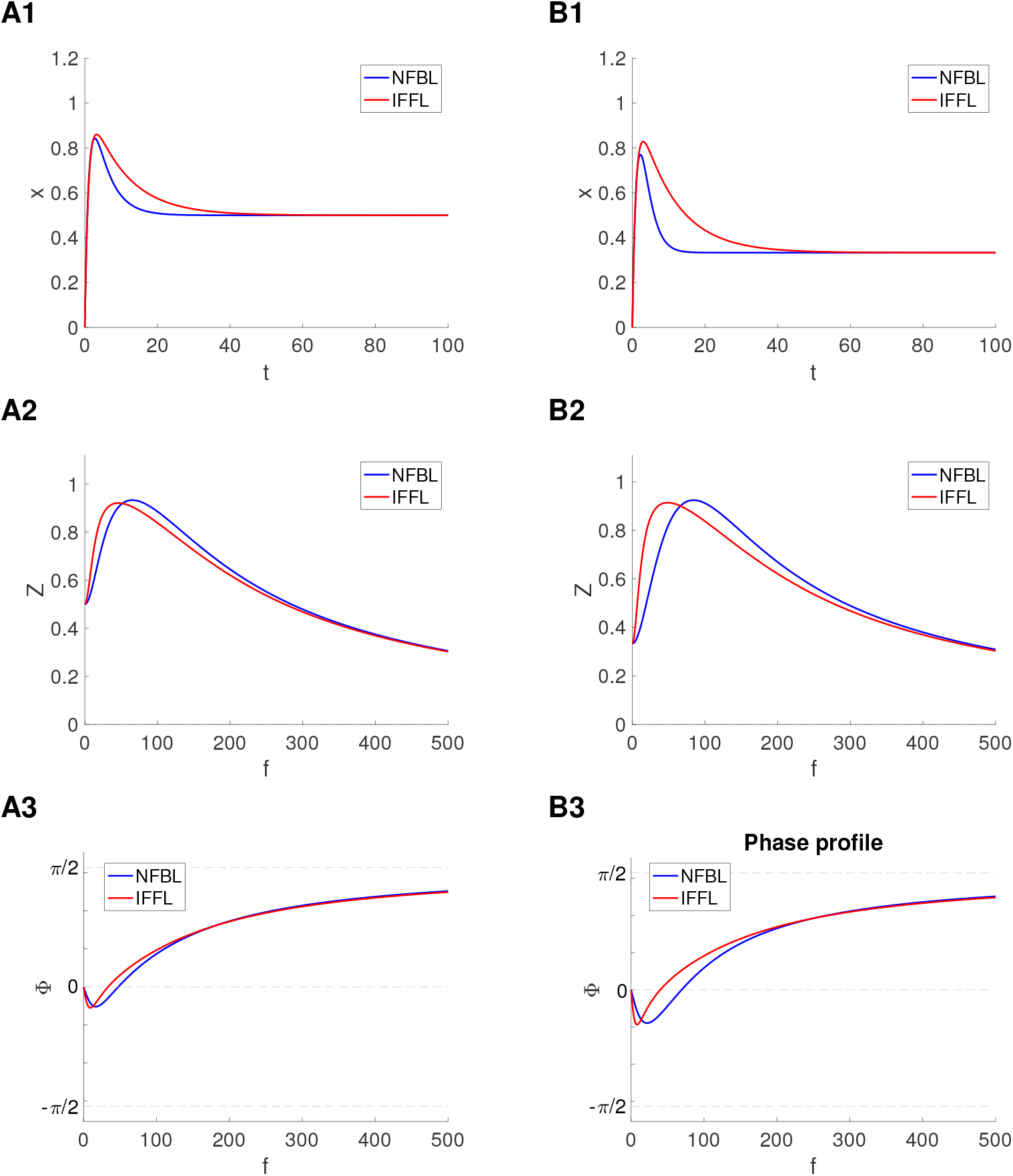
Degeneracy across circuit types is lost in non-dimensionalized linear NFBLs and IFFLs. We used System (34)-(35) with *x*(0) = 0 and *y*(0) = 0. **Top row**. Adaptive responses to constant inputs for representative parameter values for which the steady-states 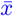 for the NFBL and IFFL coincide. **Middle row**. Impedance amplitude profiles (in response to oscillatory inputs) for the same parameter values as in panel A. **Bottom row**. Phase profiles (in response to oscillatory inputs) for the same parameter values as in panel A. In all panels *E* = 0.1. In panels B and C we used eqs. (31) and (**??**). For the model attributes we used eqs. (37). We used the following additional parameter values. **A**. NFBL: *α* = 1 and *ξ* = 0. IFFL: *α* = 0 and *ξ* = 0.5. **B**. NFBL: *α* = 2 and *ξ* = 0. IFFL: *α* = 0 and *ξ* = 0.6667.

Taken together, the results of this section show that non-dimensionalization is too stringent a reduction of a system and therefore the intra- and cross-circuit degeneracy present in the original model is hidden in the non-dimensionalized models. In contrast to the reduction method described in Section 2.2.1, non-dimensionalization involves the definition of rescaled variables in addition to the definition of the reduced parameters. From a different perspective, intra-circuit degeneracy (in the dimensional models) results from balances that keep *k*_2_*k*_3_ (NFBL) and *k*_2_*ξ* (IFFL) constant. This flexibility is lost in the non-dimensional modes since these quantities are reduced to a single parameter, which would be equivalent to making *k*_2_ = 1 in the above quantities. The existence of cross-circuit degeneracy would lead to a contradiction where *α* = 0 (*k*_3_ = 0) in the NBFL or *ζ* = 0 (*ξ* = 0) in the IFFL.

The same reasons that help identifying the general properties of the circuits and the differences between them conspire to hide the intra- and cross-circuit degeneracy since, for a given set of dimensionless parameters, the non-dimensionalized model represents a set of solutions to the original (dimensional model) spanning a large range of time scales and the other dimensional variables.

##### Remark

We note that (32) and (33) is not the only choice for dimensionless variables and parameters. For example, the choice *t*^*^ = *k*_4_ *t* leads to a different set of dimensionless variables and parameters and a dimensionless model with a different form. The results presented here remain valid.

### 3.4 Degeneracy in weakly nonlinear NFBLs and IFFLs: a window into degeneracy in nonlinear models

In the previous sections we discussed the presence of degeneracy within and across NFBLs and IFFLs in linear models and we showed that these types of degeneracy cannot be resolved by time-dependent inputs within a relatively large class. The question arises of whether and under what conditions this persists for nonlinear NFBLs and IFFLs.

In principle, there is no reason to expect degeneracy to be communicated from linear to nonlinear systems. The types of nonlinearities we consider in this paper (see Section 2.1.2) involve the (hidden) variable *y*, which, as we showed in Section 3.1.3, has a different degeneracy structure than the (observable) variable *x* in linear NFBLs and IFFLs. This is particularly relevant, for example, for the model (3)-(4) with a nonlinearity of the form (6) that involves the product of *x* and *y*.

Here we begin to examine these ideas by using weakly nonlinear models. Nonlinear systems cannot be solved analytically and therefore one typically relies on numerical simulations. Weakly nonlinear systems can be thought of as an intermediate step that provides a window into the dynamics of nonlinear systems by using linear tools and asymptotic expansions. We show that weakly nonlinear NFBLs and IFFLs inherit the intra-circuit degeneracy of the linear systems to the first order approximation, but not the cross-circuit degeneracy, which breaks at the first order approximation level of analysis.

To this end, we use system (3)-(4) with a multiplicative weakly nonlinear term of the form

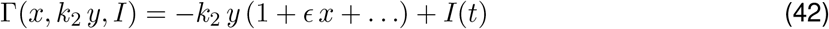

where 0 *< ϵ* ≪ 1 and *I*(*t*) is given by eq. (5).

In order to solve this system, we define

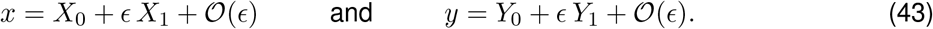

By substituting (43) into system (3)-(4), rearranging terms and collecting the terms with the same power of *E*, one obtains

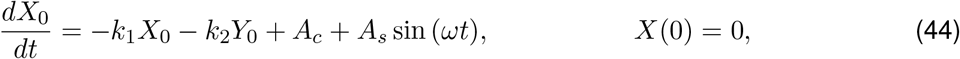

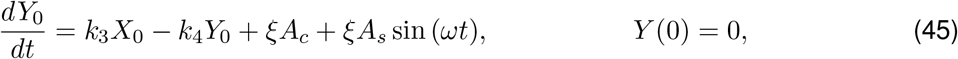

for the zeroth order approximation and

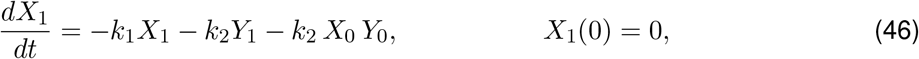

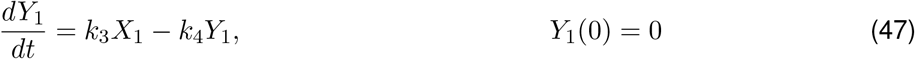

for the first order approximation.

The zeroth order approximation (44)-(45) is the linear system studied in the previous sections. Degeneracy with and across circuit types in response to constant inputs (*A*_*s*_ = 0) is as described in Section 3.1 and it persists in response to oscillatory inputs (*A*_*s*_ *>* 0) as described in Section (3.2).

Now we turn our attention to the first order approximation (46)-(47) for *A*_*s*_ = 0. This is a forced linear system where the forcing term is the product of the solutions to the zeroth order approximation. We assume that the model parameters are constrained by fixed values of the model attributes *B, C* and *Q* so that *X*_0_ exhibits degeneracy. Specifically, the parameters *k*_1_, *k*_4_ are uniquely determined by the model attributes as well as the products *k*_2_*k*_3_ (NFBL) and *k*_2_*ξ* (IFFL).

From eqs. (73)-(74), (76)-(78) and (16),

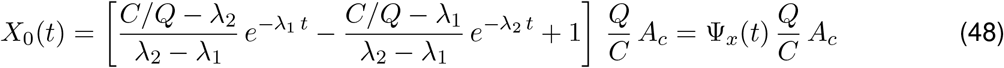

and

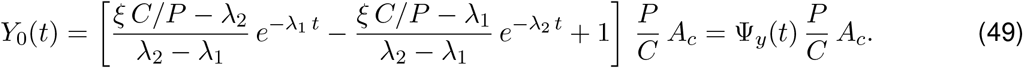

The functions Ψ_*x*_(*t*) and Ψ_*y*_(*t*) are elementary solutions to the linear system in the sense that they transition from Ψ_*x*_ = Ψ_*y*_ = 0 at *t* = 0 to Ψ_*x*_ = Ψ_*y*_ = 1 as *t* → ∞. All other solutions satisfying the linear system we are working with are multiples of these elementary functions.

For the NFBL (*ξ* = 0), *Y*_0_(*t*) simplifies to

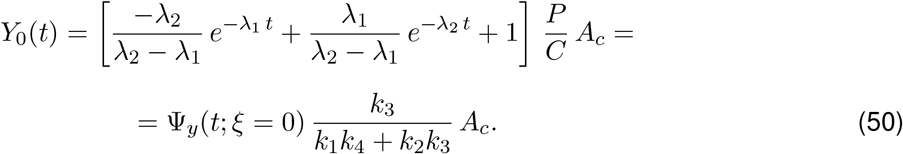

For the IFFL (*k*_3_ = 0, *C* = *k*_1_ *k*_4_, *P* = *k*_1_ *ξ*), *Y*_0_(*t*) simplifies to

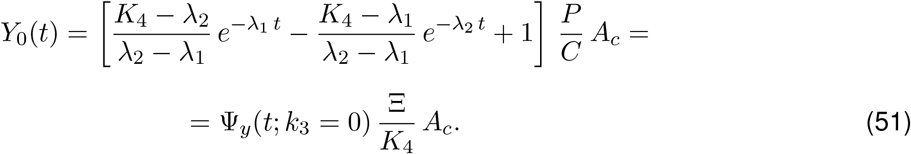

The functions Ψ_*y*_(*t*; *ξ* = 0) and Ψ_*y*_(*t*; *k*_3_ = 0) have the same degeneracy structure as *X*_0_(*t*) since they involve only the model attributes *B* and *C* (through *λ*_1_ and *λ*_2_) and *K*_4_ (IFFL), which is uniquely determined by the model attributes. Therefore, the differences between two solutions *Y*_0_(*t*) corresponding to two degenerate solutions *X*_0_(*t*) are given by factores multiplying these functions in (50) or (51). Specifically, the differences between the forcing terms −*k*_2_ *X*_0_ *Y*_0_ in (46) or (47) to two degenerate solutions *X*_0_(*t*), if they exist, are given by

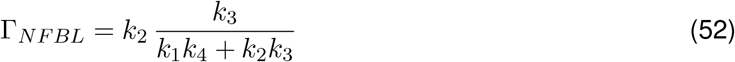

for the NFBL and

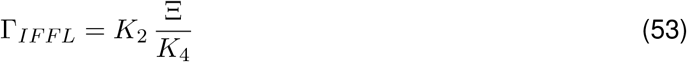

for the IFFL. Because degeneracy in *X*_0_ implies that *k*_2_ *k*_3_ (NFBL) and *K*_2_ Ξ (IFFL) are fixed, Γ_*NFBL*_ and Γ_*IFFL*_ are also fixed, accordingly. Therefore, the first order approximation to the NFBL and IFFL inherits the degeneracy (within the circuit types) from the linear zeroth order approximation.

In contrast, degeneracy across circuit types is not directly inherited from the linear zeroth order approximation. For this <u>to happe</u>n, eqs. (52) and (53) should be equal upon the transformations (*K*_1_ = 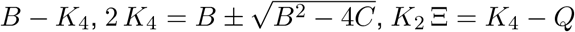 in the IFFL).

### 3.5 Degeneracy extends to nonlinear NFBLs and IFFLs with multiplicative and divisive nonlinearities

Here we examine whether the ideas discussed in the previous section persist for nonlinear NFBLs and IFFLs. To this end, we use the nonlinear models of the form (3)-(4) with a multiplicative (M-) nonlinear term given by eq. (6) and divisive nonlinear terms (Da- and Dp-) given by eqs. (7) and (8), respectively (see Section 2.1.2). The inputs are given by eq. (5). Our results are presented in Figs. 5 and 6 (M-NFBL and -IFFL), Figs. S1 and S2 (Da-NFBL and -IFFL), and Figs. S3 and S4 (Dp-NFBL and -IFFL).

**Figure 5:**
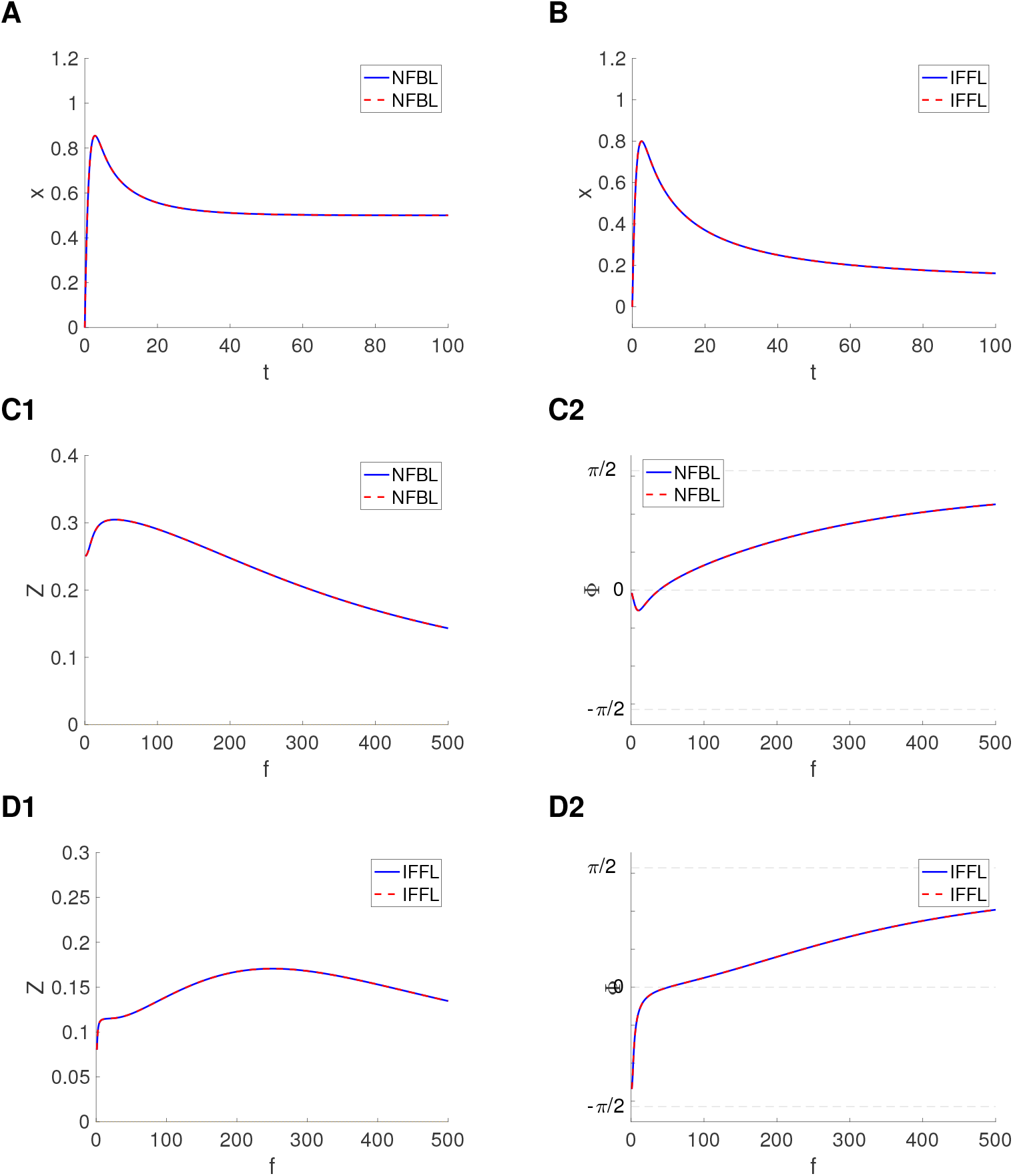
Degeneracy within circuit types in the presence of multiplicative nonlinearities: representative examples or adaptive responses (to a constant input). We used System (3)-(4) with *x*(0) = *y*(0) = 0 and Γ(*x, k*_2_ *y, I*) given by eq. (6). **A, B**. Adaptive responses for *A*_*c*_ = 1 (*A*_*s*_ = 0). **C, D**. Impedance amplitude and phase profiles for *A*_*s*_ = 1 (*A*_*c*_ = 0). **A**. Parameter values: *k*_1_ = 1, *k*_2_ = 2, *k*_3_ = 0.05, *k*_4_ = 0.05, *ξ* = 0 (blue), *k*_1_ = 1, *k*_2_ = 1, *k*_3_ = 0.1, *k*_4_ = 0.05, *ξ* = 0 (dashed-red). **B**. Parameter values: *K*_1_ = 1, *K*_2_ = 2, *K*_3_ = 0, *K*_4_ = 0.015, Ξ = 0.05 (blue), *K*_1_ = 1, *K*_2_ = 0.4, *K*_3_ = 0, *K*_4_ = 0.015, Ξ = 0.25 (dashed-red). **C**. Same parameter values and color-code as in A. **D**. Same parameter values and color-code as in B.

**Figure 6:**
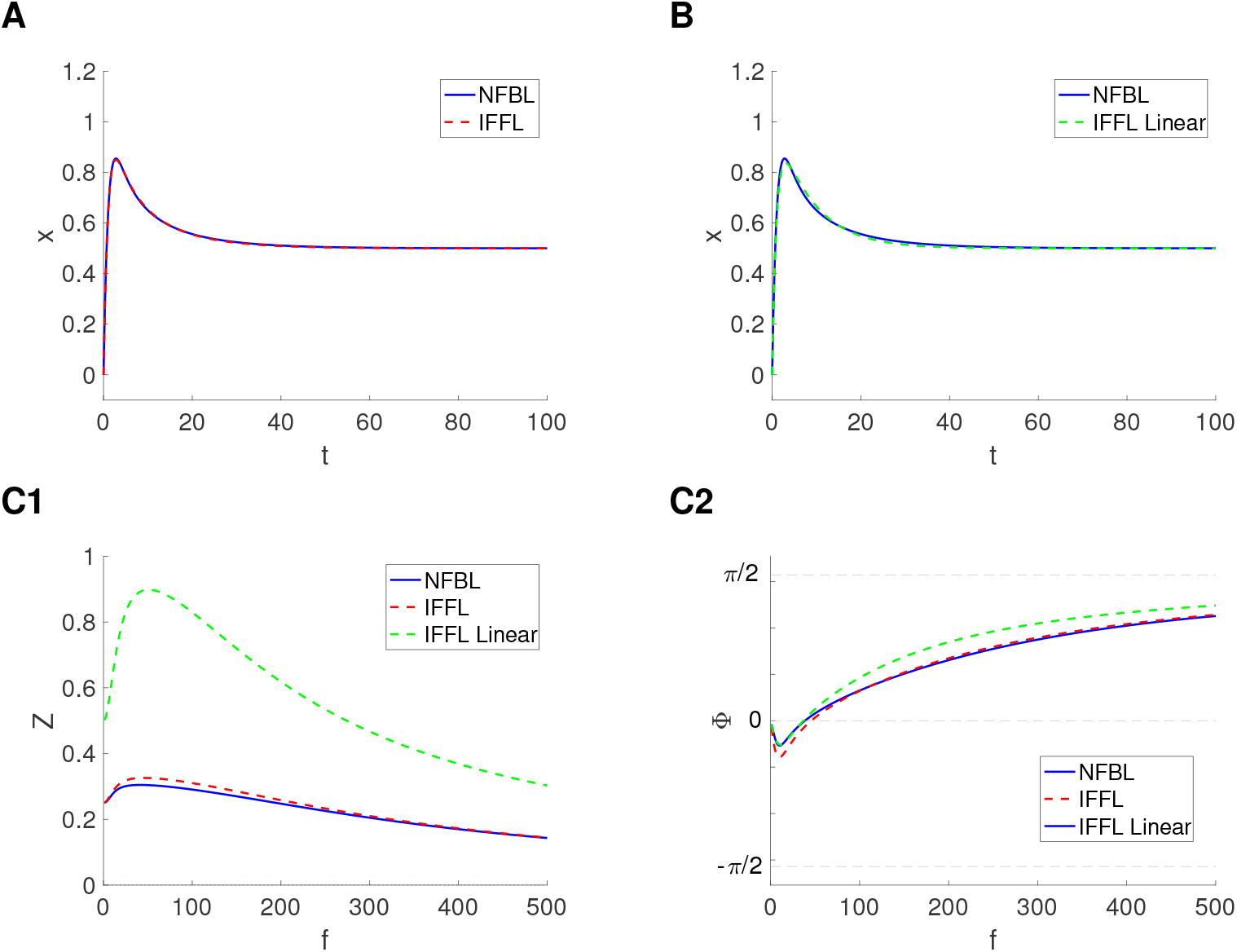
Quasi-degeneracy across circuit types in the presence of multiplicative nonlinearities: representative examples of adaptive responses (to a constant input). We used System (3)-(4) with *x*(0) = *y*(0) = 0 and Γ(*x, k*_2_ *y, I*) given by (6). **A, B**. Adaptive responses for *A*_*c*_ = 1 (*A*_*s*_ = 0). **C**. Impedance amplitude and phase profiles for *A*_*s*_ = 1 (*A*_*c*_ = 0). Degeneracy is disrupted in response to oscillatory inputs. **A**. Parameter values: *k*_1_ = 1, *k*_2_ = 2, *k*_3_ = 0.05, *k*_4_ = 0.05, *ξ* = 0 (blue), *K*_1_ = 0.9525, *K*_2_ = 2, *K*_3_ = 0, *K*_4_ = 0.086, Ξ = 0.045 (dashed-red). **B**. Parameter values: *k*_1_ = 1, *k*_2_ = 2, *k*_3_ = 0.05, *k*_4_ = 0.05, *ξ* = 0 (blue). For the linear model we used System (1)-(2) with *K*_1_ = 1.0102, *K*_2_ = 2, *K*_3_ = 0, *K*_4_ = 0.1216, Ξ = 0.0301. For these parameter values, the eigenvalues are given by *r*_1_ = −0.1216 and *r*_2_ = −1.0102. **C**. Quasi-degeneracy breaks in response to oscillatory inputs. The parameter values and color-code are as in A and B.

**Figure 7:**
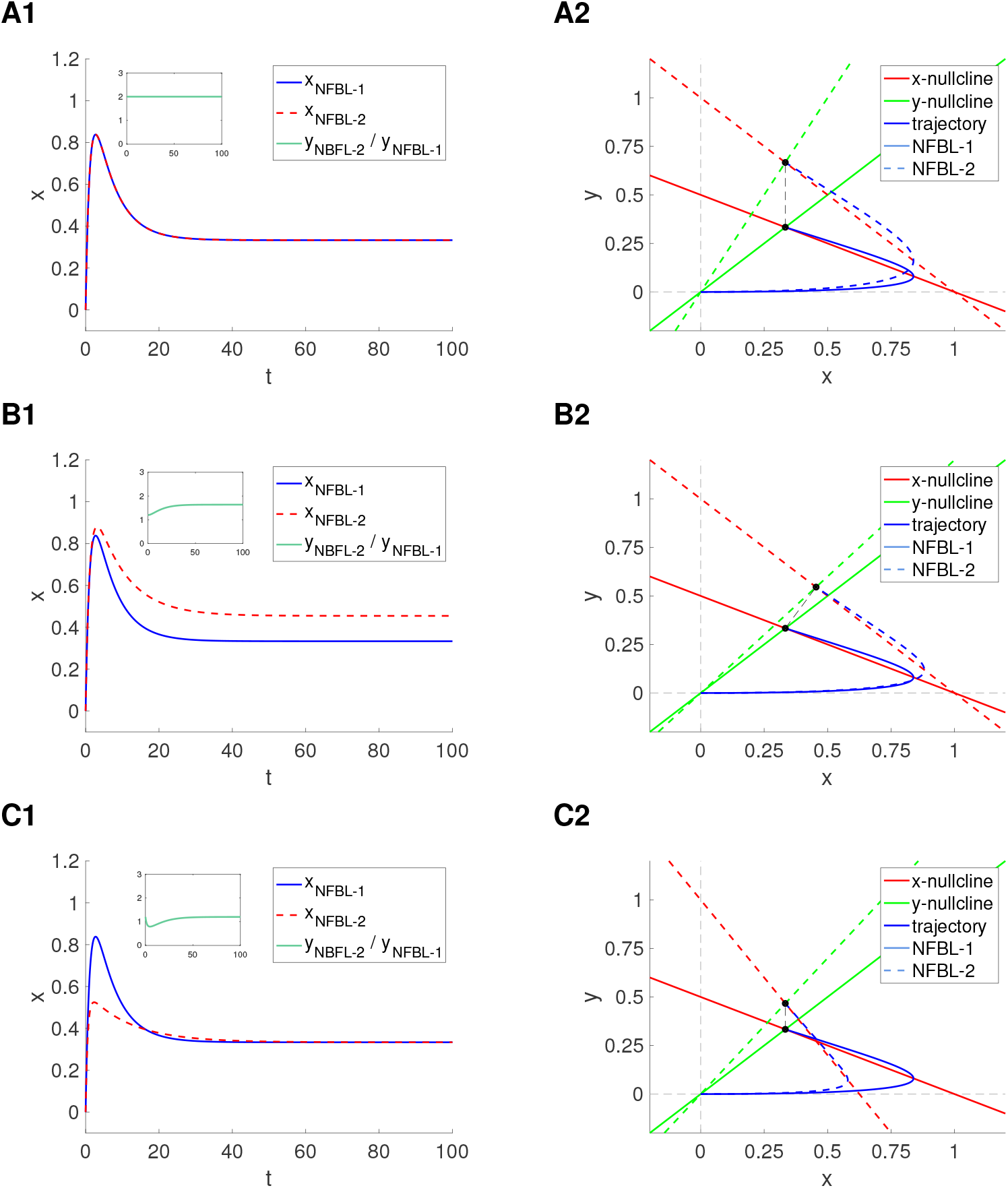
Degeneracy of adaptive patterns (responses to constant inputs) : Phase-plane analysis. We used system (1)-(2) with *A*_*c*_ = 1, *A*_*s*_ = 0, *x*(0) = 0 and *y*(0) = 0. **Left column**. Superimposed *x*-traces for two NFBLs: NFBL-1 (blue) and NFBL-2 (dashed-red). Inset: quotient of corresponding *y*-traces. **Right column**. Superimposed phase-plane diagrams for the two cases in the corresponding left column. The *x*-nullclines (red), *y*-nullclines (green) and trajectories (blue) are presented with solid lines (NBFL-1) and dashed lines (NFBL-2). The two fixed-points are indicated with black dots. The dashed-black line connects between them. **A**. Degeneracy within NFBLs. Parameter values: *k*_1_ = 1, *k*_2_ = 2, *k*_3_ = 0.05, *k*_4_ = 0.05 (NFBL-1, blue), *k*_1_ = 1, *k*_2_ = 1, *k*_3_ = 0.1, *k*_4_ = 0.05 (NBFL-2, dashed-red). **B**. No degeneracy within NFBLs. The adaptive patterns have different steady-state values (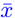). Parameter values: *k*_1_ = 1, *k*_2_ = 2, *k*_3_ = 0.05, *k*_4_ = 0.05 (NFBL-1, blue), *k*_1_ = 1, *k*_2_ = 1, *k*_3_ = 0.06, *k*_4_ = 0.05 (NBFL-2, dashed-red). **C**. No degeneracy within NFBLs. The adaptive patterns have the same steady-state value (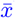). Parameter values: *k*_1_ = 1, *k*_2_ = 2, *k*_3_ = 0.05, *k*_4_ = 0.05 (NFBL-1, blue), *k*_1_ = 1.6, *k*_2_ = 1, *k*_3_ = 0.07, *k*_4_ = 0.05 (NBFL-2, dashed-red).

#### 3.5.1 Intra-circuit-type degeneracy is present in NFBLs and IFFLs in response to constant inputs

Figs. 5-A and -B illustrate the presence of (intra-circuit-type) degeneracy in the adaptive patterns produced by the M-NFBL (Figs. 5-A) and M-IFFL (Figs. 5-B) for two different combinations of *k*_2_ and *k*_3_ (NFBL) and *K*_2_ and Ξ (IFFL), thus extending our results for linear and weakly nonlinear NFBLs and IFFLs discussed above.

While it is not surprising that balances between two opposing processes (inhibition of node X by node Y and either activation of the node Y by node X or direct activation of the node Y by the input; Fig. 1-A) lead to the same steady-state response (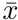), it is not a priori clear that they would necessarily lead to degeneracy in nonlinear models in both circuit types, which involve dynamic processes. The two divisive models show similar results (Figs. S1-A and -B for the Da-NFBL and -IFFL and Figs. S3-A and -B the Dp-NFBL and -IFFL).

#### 3.5.2 Intra-circuit-type degeneracy persists in NFBLs and IFFLs in response to oscillatory inputs

We measured the model’s response to oscillatory inputs of the from (5) by using the definitions of *Z* and Φ given by eqs. (24) (see Section 2.3.2).

Figs. 5-C and -D illustrate that the degeneracy structure in response to oscillatory signals is identical to the degeneracy structure in response to constant inputs. These Figs. show the impedance (Figs. 5-C1 and -D1) and phase (Figs. 5-C2 and -D2) profiles for the M-NFBL ((Fig. 5-C) and M-IFFL (Fig. 5-D) for the same set of parameter values as in Figs. 5-A1 and -B1. These results indicate that, similarly to the corresponding linear models, degeneracy within NFBLs and IFFLs cannot be resolved by oscillatory signals, and more generally time-dependent signals within a large class. The divisive models exhibit similar behavior (Figs. S1-C and -D for the Da-NFBL and -IFFL and Figs. S3-C and -D the Dp-NFBL and -IFFL).

#### 3.5.3 Quasi-degeneracy across NFBLs and IFFLs in response to constant inputs

Uncovering degeneracy across linear NFBLs and IFFLs was mathematically facilitated by the formulas relating the model parameters with the model attributes (*B, C* and *Q*). In contrast to the intra-circuit-type degeneracy where degeneracy reflects different possible balances between the activation of *y* (*k*_3_ or *ξ*) and the inhibition it exerts on *x* (*k*_2_), degeneracy across circuit types reflects more complex relationships between the participating parameters.

Since we have no access to analytic formulas for nonlinear models, we searched for solutions to the M-IFFL model that approximate the previously computed solutions to the M-NFBL model (Figs. 5-A, blue). To this end, we used a gradient descent algorithm to compute the optimal parameter values for *K*_1_ and *K*_4_ (within an error of 10^−6^). The superimposed adaptive patterns are shown in Fig. 6-A. We refer to this phenomenon as quasi-degeneracy since the solutions are not mathematically identical, although almost indistinguishable for practical purposes. Additional quasi-degenerate adaptive patterns are provided by the multiple combinations of *K*_2_ and Ξ that produce degeneracy within the M-IFFL (as discussed in the previous section). This type of quasi-degenerate adaptive patterns are also obtained for the Da- and Dp-NFBL and -IFFL models (Figs. S2-A and S4-A).

#### 3.5.4 Quasi-degeneracy across NFBLs and IFFLs and across linear and nonlinear circuits in response to constant inputs

Because the adaptive patterns for the nonlinear M-NFBL and -IFFL resemble the adaptive patterns for the linear NFBL and IFFL, we asked whether there exists quasi-degeneracy both across circuit types and across linear and nonlinear models. To this end used a gradient descent algorithm to compute the optimal values of *λ*_1_ and *λ*_2_ in the analytical solution to linear models (see Section B.1) that approximates the solution to the M-NFBL (within an error of 10^−6^). (We used the same parameter values as in Figs. 5-A and 6-A, blue curves.) We then reconstructed the linear IFFL model (see Supplementary Material) from this optimal solution by recovering the parameter values. The superimposed adaptive patterns are shown in Fig. 6-B. Additional quasi-degenerate adaptive patterns are provided by the (intra-circuit-type) degenerate adaptive patterns for linear IFFLs discussed above (not shown).

We repeated the process for the Da-NFBL model (for the same parameter values as in Figs. S1-A and S2-A, blue curves.) We computed the quasi-degenerate, optimal solution for linear models (within an error of 10^−6^). Our result is shown in Fig. S1-B. The parameter *η* in *C*_1_ and *C*_2_ is given by *η* = −*k*_1_*x*(0)+1*/*(1+*k*2*y*(0))+*A* = 1+*A* instead of *η* = −*k*_1_*x*(0) −*k*_2_*y*(0)+*A* = *A* as for the M-IFFL. This has consequences for the reconstruction of the Da-IFFL model since the appropriate initial condition for the linear model is *y*(0) = − (*η*− *A*)*/K*_2_, which is different from the initial condition for the Da-NFBL and -IFFL models, *y*(0) = 0. Additionally, for the linear model, the recovered parameter Ξ *<* 0. For the same recovered parameter values, but *y*(0) = 0, the solution not only show no adaptation, but it is monotonically increasing. In other words, the adaptive patterns in the reconstructed linear model are generated by the initial value of the variable *y* that causes *x* to increases and an inhibitory input signal that causes *x* to peak and subsequently decrease. This mechanism is different from the M- and Da-NFBL models the linear IFFL model is approximating. We also repeated the process for the Dp-NFBL model and obtained results similar to the M-NFBL model (Fig. S4-B).

#### 3.5.5 Quasi-degeneracy across NFBLs and IFFLs is disrupted in response to oscillatory inputs

Fig. 6-C shows the impedance (Fig. 6-C1) and phase (Fig. 6-C2) profiles for the M-NFBL and -IFFL models discussed above (using the same parameter values as in Figs. Fig. 6-A and -B). The small difference between the adaptive patterns for the M-NFBL and M-IFFL are reflected in larger difference between the corresponding impedance and phase profiles (compare the blue and dashed-red curves). Similarly, the difference between the adaptive patterns for the nonlinear NFBL and the linear IFFL are reflected in larger differences between the corresponding impedance and phase profiles (compare the blue and dashed-green curves). Interestingly, the differences between the impedance and phase profiles between the linear IFFL and the nonlinear NFBL and IFFL are significantly larger than the differences between the corresponding adaptive patterns. In other worlds, oscillatory inputs and more generally time-dependent inputs are potentially useful tools to resolve the quasi-degeneracy present in the adaptive patterns by amplifying the small differences between them.

Fig. S2-C shows the impedance (Fig. S2-C1) and phase (Fig. S2-C2) profiles for the Da-NFBL and -IFFL models discussed above (using the same parameter values as in Figs. Fig. S2-A and -B). The results for the nonlinear models are analogous to the ones described above: a small difference between the adaptive patterns for the Da-NFBL and Da-IFFL are reflected in larger difference between the corresponding impedance and phase profiles (compare the blue and dashed-red curves). In contrast, the impedance and phase profiles for the linear IFFL model is qualitatively different from the other two. Specifically, the IFFL exhibits a low-pass filter (no resonance) and a monotonically increasing phase (no phasonance). This is a direct consequence of the response to constant input being a monotonically increasing function for zero initial conditions as opposed to an adaptive pattern.

The results for the Dp-NFBL and -IFFL models are similar to the corresponding M-models (Figs. S4-C and -D).

In summary, quasi-degeneracy across circuit types for nonlinear NFBLs and IFFLs can be resolved by oscillatory inputs and, more generally, by time-dependent inputs.

### 3.6 A dynamical systems mechanistic interpretation of degeneracy in adaptive NFBLs and IFFLs

Here we use dynamical system tools to analyze the mechanisms of generation of adaptive patterns (in response to constant inputs) within and across circuit types. We use system (1)-(2) with *A*_*c*_ = 1, *A*_*s*_ = 0, *x*(0) = 0 and *y*(0) = 0.

#### 3.6.1 Dynamic balances responsible for the generation of degeneracy within NFBLs

Fig. 7-A1 shows an example of degenerate adaptive patterns for *k*_2_ = 2, *k*_3_ = 0.05 (NFBL-1, blue), *k*_2_ = 1, *k*_3_ = 0.1 (NBFL-2, dashed-red) and the same values of *k*_1_ and *k*_4_. (The inset shows that the quotient of the corresponding *y*-traces is constant, see Section B.2). Fig. 7-A2 shows the superimposed phase-plane diagrams for the two parameter sets in Fig. 7-A1 (solid curves for NFBL-1 and dashed curves for NFBL-2). The *x*- and *y*-nullclines are given, respectively, by

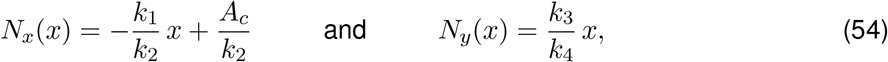

and the fixed-points (computed as the intersection of the two nullclines) s given by

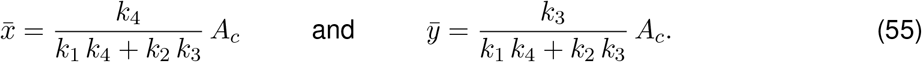

We first describe the geometry of the generation of adaptive patterns by focusing on NFBL-1 (blue curve in Fig. 7-A1 and solid curves in Fig. 7-A2). The adaptive trajectory transitions from (0, 0) to (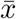, 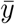) (black dot). The overshoot is created because the structure of the vector field requires the trajectory to first move to the right (increasing values of *x*), then cross the *x*-nullcline (red), thus generating the peak of the adaptive pattern, and finally move to the left (decreasing values of *x*) towards the fixed-point.

The transition from NFBL-1 to NFBL-2 consists of a decrease in *k*_2_ (inhibition) and an increase in *k*_3_ (activation). While 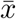 remains constant (necessary condition for degeneracy), 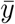 increases (Fig. 7-A2, black dots). The quotient between the two values of *k*_3_ (0.1*/*0.05 = 2) determines the value of the constant quotient of the two *y*-traces (see Section B.2) (Fig. 7-A1, inset).

The decrease in *k*_2_ and the increase in *k*_3_ cause the slopes of the *x*- and *y*-nullclines, respectively, to increase. This in turn causes the fixed-point to raise. These changes need to be balanced so the two trajectories cross the corresponding *x*-nullclines for the same value of *x* and evolve towards the fixed-point by maintaining a constant quotient of the variables *y*. In other words, if the *x*-nullcline’s slope increases (due to a decrease in *k*_2_), then the *y*-nullcline’s slope needs to increase (by increasing *k*_3_) in such a way as to maintain the constancy of the quotient of the variables *y*. Conversely, if the *y*-nullcline’s slope increases (due to an increase in *k*_3_, then the *x*-nullcline’s slope needs to increase (by decreasing *k*_2_) in such a way as to maintain the constancy of the quotient of the variables *y*. In order for the trajectories to cross the *x*-nullclines for the same values of *x*, the two *x*-nullclines need to intersect for a values of *x* higher than the crossing value (peak of the adaptive pattern).

If the *y*-nullcline’s slope does not increase enough (Fig. 7-B2, (*k*_2_, *k*_3_) transitions from (2, 0.05) for NFBL-1 to (1, 0.06) for NFBL-2), then the two fixed-points are not on a vertical line and degeneracy is broken (Fig. 7-B1).

Maintaining the two fixed-points on a vertical line does not require *k*_2_ *k*_3_ to be fixed for both NFBL-1 and NFBL-2 (Fig. 7-C2, (*k*_2_, *k*_3_) transitions from (2, 0.05) for NFBL-1 to (1, 0.07) for NFBL-2). However, this requires *k*_1_ to change as well (*k*_1_ = 1 for NBFL-1 and *k*_1_ = 1.6 for NFBL-2). This causes the *x*-nullcline’s slope to increase, but the *x*-nullcline does not simply shifts upwards (by maintaining the same *x*-intercept), but it crosses the other *x*-nullcline at a non-zero values of *x* (the two *x*-nullclines have different *x*-intercepts). As a result, the trajectory for NFBL-2 crosses the corresponding *x*-nullcline at a lower values of *x* than the trajectory for NBFL-1, and therefore degeneracy is broken (Fig. 7-C1). Note the quotient of the *y* variables is not constant (inset).

#### 3.6.2 Dynamic balances responsible for the generation of degeneracy within IFFLs

Fig. 8-A1 shows an example of degenerate adaptive patterns for *k*_2_ = 2, *ξ* = 0.015 (IFFL-1, blue), *k*_2_ = 1, *ξ* = 0.03 (IFFL-2, dashed red) and the same values of *k*_1_ and *k*_4_. (The inset shows that the quotient of the corresponding *y*-traces is constant, see Section B.2). Fig. 8-A2 shows the superimposed phase-plane diagrams for the two parameter sets in Fig. 8-A1(solid curves for IFFL-1 and dashed curves for IFFL-2). The *x*-nullclines are given by eq. (54) (left) and the *y*-nullcline is given by

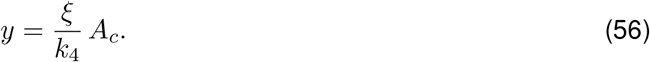

**Figure 8:**
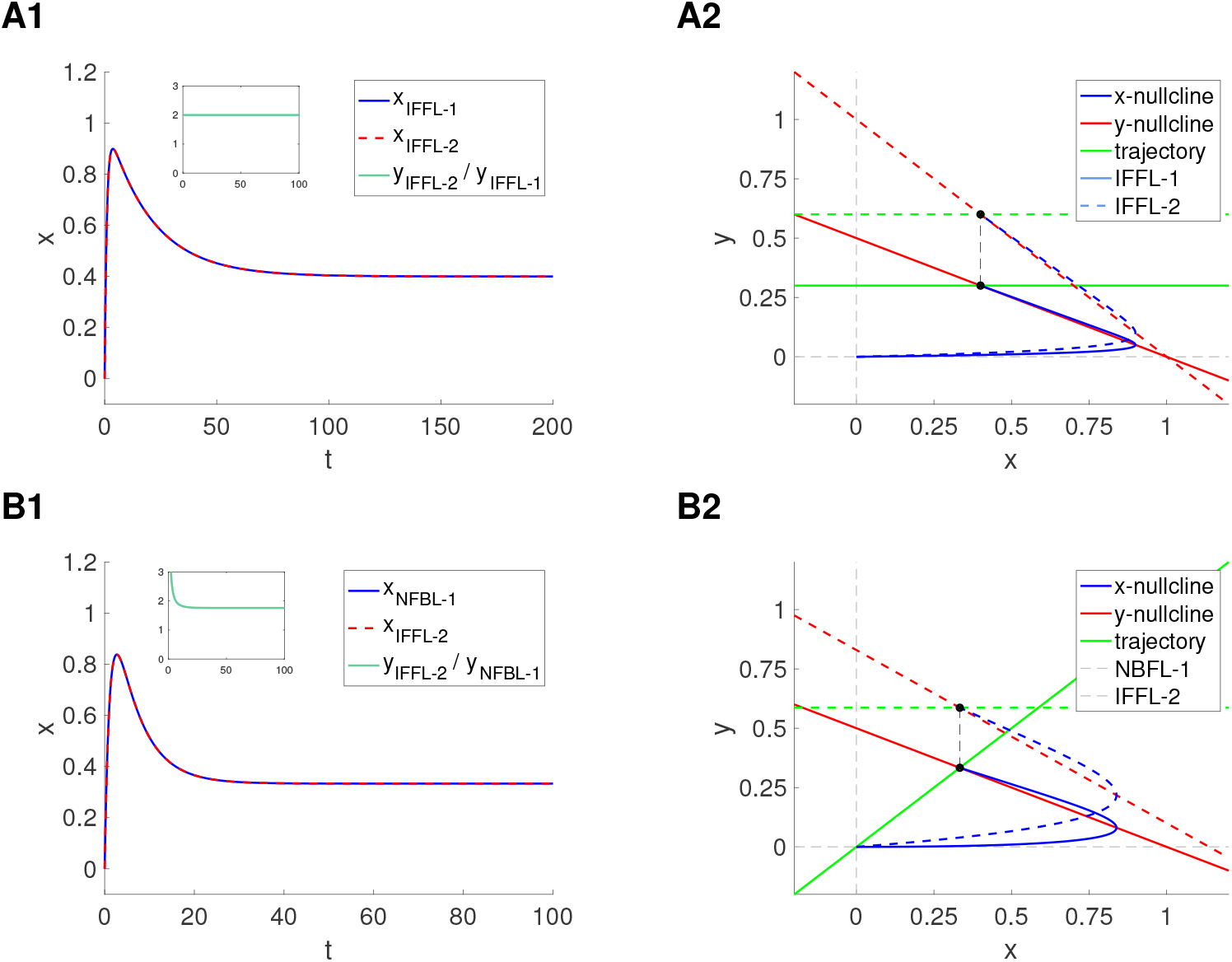
Degeneracy of adaptive patterns (responses to constant inputs): Phase-plane analysis. We used system (1)-(2) with *A*_*c*_ = 1, *A*_*s*_ = 0, *x*(0) = 0 and *y*(0) = 0. **Left column**. Superimposed *x*-traces for two NFBLs: NFBL-1 (blue) and NFBL-2 (dashed-red). Inset: quotient of corresponding *y*-traces. **Right column**. Superimposed phase-plane diagrams for the two cases in the corresponding left column. The *x*-nullclines (red), *y*-nullclines (green) and trajectories (blue) are presented with solid lines (NBFL-1) and dashed lines (NFBL-2). The two fixed-points are indicated with black dots. The dashed-black line connects between them. **A**. Degeneracy within IFFLs. Parameter values: *k*_1_ = 1, *k*_2_ = 2, *k*_4_ = 0.05, *ξ* = 0.015 (IFFL-1, blue), *k*_1_ = 1, *k*_2_ = 1, *k*_4_ = 0.05, Ξ = 0.03 (IFFL-2, dashed-red). **B**. Degeneracy across NFBL and IFFL. Parameter values: *k*_1_ = 1, *k*_2_ = 2, *k*_3_ = 0.05, *k*_4_ = 0.05 (NFBL-1, blue), *k*_1_ = 0.8794, *k*_2_ = 1.2056, *k*_4_ = 0.1706 and *ξ* = 0.1.

The fixed-points are given by

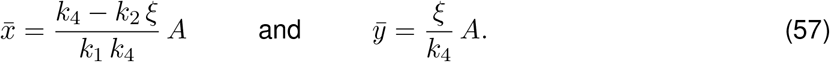

The mechanism of generation of adaptive patterns is similar to the mechanism described above. Briefly, for the two degenerate adaptive patterns, *k*_2_ Ξ = 0.03. The transition from IFFL-1 to IFFL-2 consists of a decrease in *k*_2_ (inhibition) and an increase in *ξ* (direct activation). While 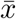 remains constant (necessary condition for degeneracy), 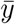 increases (Fig. 8-A2, black dots). The quotient between the two values of *ξ* (0.1*/*0.05 = 2) determines the value of the constant quotient of the two *y*-traces (see Section B.2) (Fig. 8-A1, inset).

The decrease in *k*_2_ and the increase in *ξ* cause the slope of the *x*-nullcline to increase and the *y*-nullcline to raise, respectively.This in turn causes the fixed-point to raise. These changes need to be balanced so the two trajectories cross the corresponding *x*-nullclines for the same value of *x* and evolve towards the fixed-point by maintaining a constant quotient of the variables *y*. In other words, if the *x*-nullcline’s slope increases (due to a decrease in *k*_2_), then the *y*-nullcline needs to raise (by increasing *ξ*) in such a way as to maintain the constancy of the quotient of the variables *y*. In order for the trajectories to cross the *x*-nullclines for the same values of *x*, the two *x*-nullclines need to intersect for a values of *x* higher than the crossing value (peak of the adaptive pattern).

#### 3.6.3 Dynamic balances responsible for the generation of degeneracy across NFBLs and IFFLs

Fig. 8-B1 shows an example of degenerate adaptive patterns for *k*_1_ = 1, *k*_2_ = 2, *k*_3_ = 0.05, *k*_4_ = 0.05 (NFBL-1, blue), *k*_1_ = 0.8794, *k*_2_ = 1.2056, *k*_4_ = 0.1706 and *ξ* = 0.1 (IFFL-2, dashed-red). Fig. 8-B2 shows the superimposed phase-plane diagrams for the two parameter sets in Fig. 8-B1 (solid curves for NFBL-1 and dashed curves for IFFL-2). Together with the ideas discussed above, they illustrate the mechanism of generation of degenerate adaptive patterns is a combination of the ideas discussed in the previous section.

#### 3.6.4 The dynamic balances responsible for the generation of degeneracy within and across nonlinear NFBLs and IFFLs are inherited from the linear regimes

Here we extend our investigation of the mechanism of generation of degeneracy within and across NFBLs and IFFLs to include nonlinear models (3)-(4) with multiplicative (6) and divisive (7) nonlinearities. Our results are presented in Fig. 9 (multiplicative) and Fig. 10 (divisive). In both cases, the nonlinearities affect the *x*-nullclines, but not the *y*-nullclines. The latter are given by eq. (54) (NFBL, right) and eq. (56) (IFFL). The *x*-nullclines for the multiplicative and divisive models are given, respectively, by

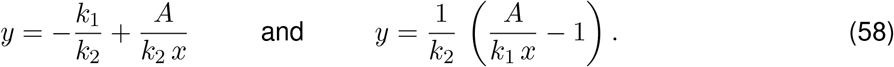

**Figure 9:**
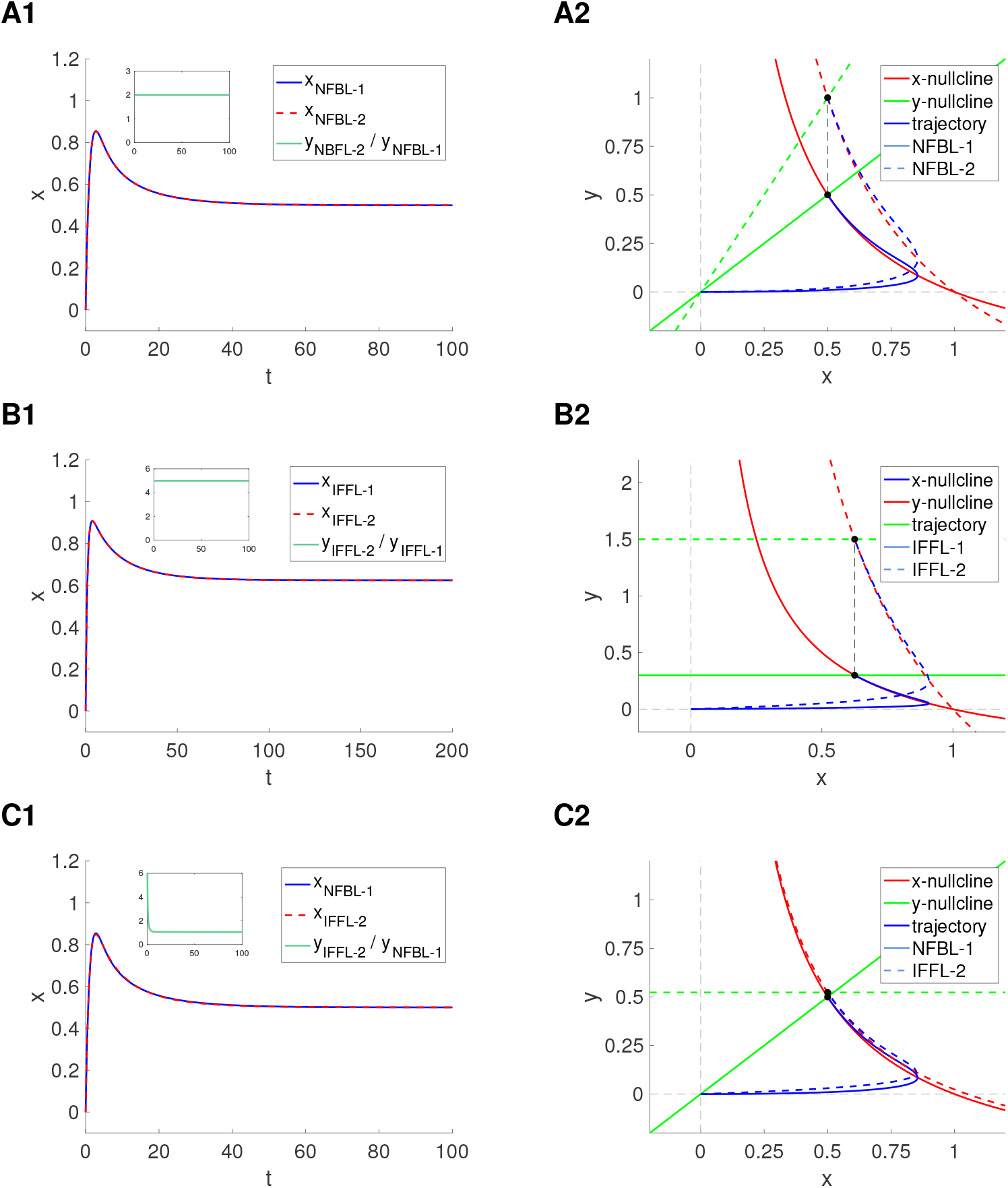
Degeneracy of adaptive patterns (responses to constant inputs) in nonlinear NFBLs and IFFLs with multiplicative nonlinearities: Phase-plane analysis. We used System (3)-(4) with *x*(0) = *y*(0) = 0 and Γ(*x, k*_2_ *y, I*) given by eq. (6). **Left column**. Superimposed *x*-traces for two NFBLs: NFBL-1 (blue) and NFBL-2 (dashed-red). Inset: quotient of corresponding *y*-traces. **Right column**. Superimposed phase-plane diagrams for the two cases in the corresponding left column. The *x*-nullclines (red), *y*-nullclines (green) and trajectories (blue) are presented with solid lines (NBFL-1) and dashed lines (NFBL-2). The two fixed-points are indicated with black dots. The dashed-black line connects between them. **A**. Degeneracy within NFBLs. Parameter values: *k*_1_ = 1, *k*_2_ = 2, *k*_3_ = 0.05, *k*_4_ = 0.05, *ξ* = 0 (blue), *k*_1_ = 1, *k*_2_ = 1, *k*_3_ = 0.1, *k*_4_ = 0.05, *ξ* = 0 (dashed-red). **B**. Degeneracy within IFFLs. Parameter values: *k*_1_ = 1, *k*_2_ = 2, *k*_3_ = 0, *k*_4_ = 0.05, *ξ* = 0.015 (blue), *k*_1_ = 1, *k*_2_ = 0.4, *k*_3_ = 0, *k*_4_ = 0.05, *ξ* = 0.075 (dashed-red). **C**. Quasi-degeneracy across NFBLs and IFFLs. Parameter values: *k*_1_ = 1, *k*_2_ = 2, *k*_3_ = 0.05, *k*_4_ = 0.05, *ξ* = 0 (blue) *k*_1_ = 0.9525, *k*_2_ = 2, *k*_3_ = 0, *k*_4_ = 0.086, *ξ* = 0.045 (dashed-red).

**Figure 10:**
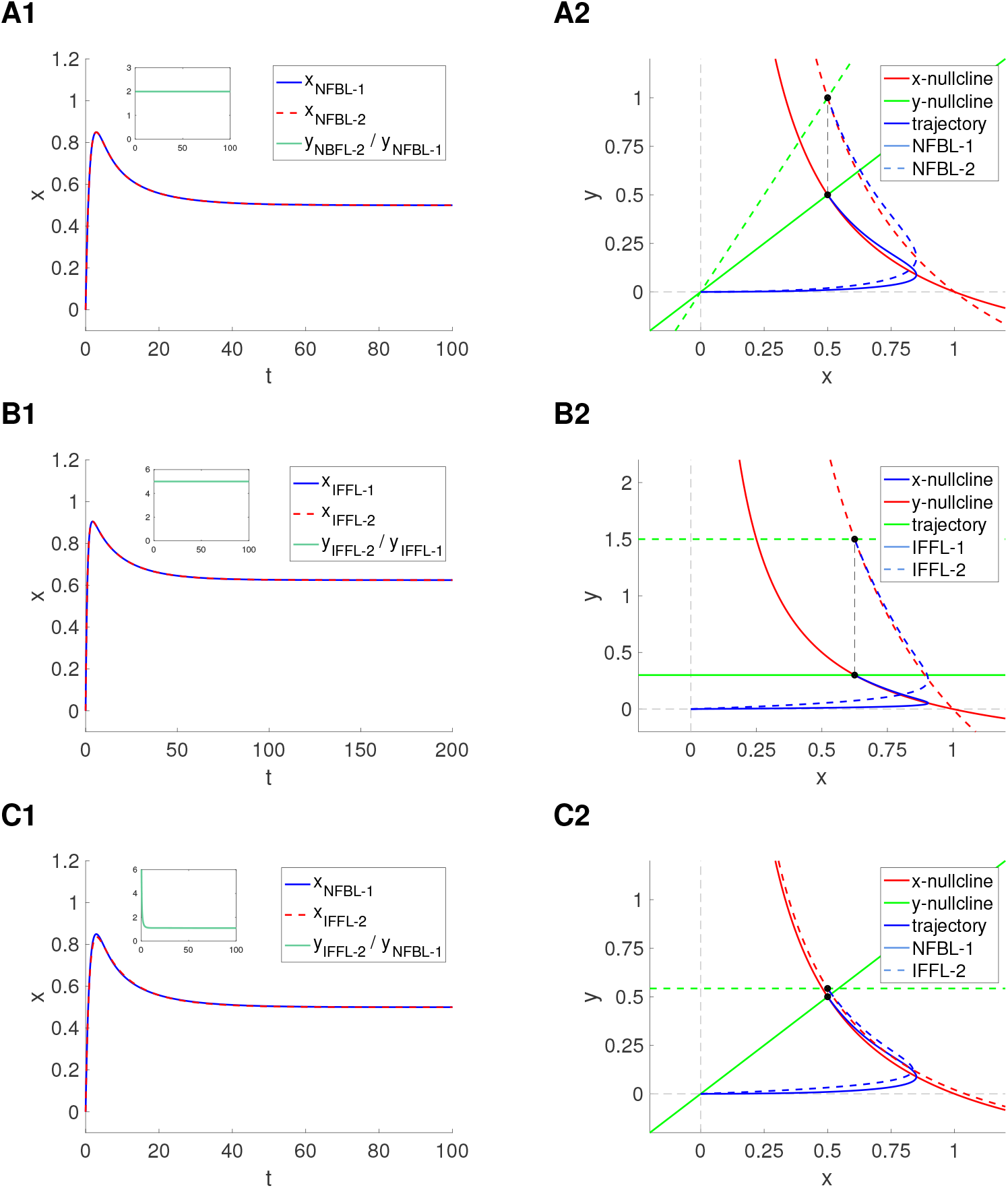
Degeneracy of adaptive patterns (responses to constant inputs) in NFBLs and IFFLs with divisive nonlinearities: Phase-plane analysis. We used System (3)-(4) with *x*(0) = *y*(0) = 0 and Γ(*x, k*_2_ *y, I*) given by eq. (7). **Left column**. Superimposed *x*-traces for two NFBLs: NFBL-1 (blue) and NFBL-2 (dashed-red). Inset: quotient of corresponding *y*-traces. **Right column**. Superimposed phase-plane diagrams for the two cases in the corresponding left column. The *x*-nullclines (red), *y*-nullclines (green) and trajectories (blue) are presented with solid lines (NBFL-1) and dashed lines (NFBL-2). The two fixed-points are indicated with black dots. The dashed-black line connects between them. **A**. Degeneracy within NFBLs. Parameter values: *k*_1_ = 1, *k*_2_ = 2, *k*_3_ = 0.05, *k*_4_ = 0.05, *ξ* = 0 (blue), *k*_1_ = 1, *k*_2_ = 1, *k*_3_ = 0.1, *k*_4_ = 0.05, *ξ* = 0 (dashed-red). **B**. Degeneracy within IFFLs. Parameter values: *k*_1_ = 1, *k*_2_ = 2, *k*_3_ = 0, *k*_4_ = 0.05, *ξ* = 0.015 (blue), *k*_1_ = 1, *k*_2_ = 0.4, *k*_3_ = 0, *k*_4_ = 0.05, *ξ* = 0.075 (dashed-red). **C**. Quasi-degeneracy across NFBLs and IFFLs. Parameter values: *k*_1_ = 1, *k*_2_ = 2, *k*_3_ = 0.05, *k*_4_ = 0.05, *ξ* = 0 (blue) *k*_1_ = 0.9594, *k*_2_ = 2, *k*_3_ = 0, *k*_4_ = 0.0855, *ξ* = 0.0464 (dashed-red).

The fixed-points are computed as the intersection between the corresponding *x*- and *y*-nullclines. Note that these nullclines are identical for *k*_1_ = 1.

The phase-plane diagramas in Figs. 9 and 10 are different from the corresponding ones for the linear models (Figs. 7-A and 6). However, their structures are qualitatively similar. In fact, geometrically, the phase-plane diagrams for the nonlinear models can be thought of as deformation of the phase-plane diagrams for the corresponding linear models. Therefore, the mechanism responsible for the generation of degeneracy in the nonlinear multiplicative and divisive models can be explained following the same ideas as for the linear models discussed above. This explains why one can relatively easily find quasi-degenerate adaptive patterns between linear and nonlinear models. One consequence of this is that the mechanisms of generation of degeneracy are qualitatively similar between the multiplicative and divisive models. This is supported by the fact that although the two models are qualitatively different, the phase-plane diagrams are very similar (in some cases the *x*-nullclines are identical as mentioned above).

## 4 Discussion

Biological adaptation has received significant attention in recent years [1–6]. The identification of NFBLs and IFFLs as two elementary building block circuits able to produce these types of adaptive patterns [3] motivated the investigation of methods to discriminate between these two circuit types [4,26–29]. These efforts require an understanding of the biological and dynamic mechanisms that give rise to adaptive patterns (in response to step-constant inputs) and, more generally, how network processing (NFBL) and input orchestration (IFFL) control the responses of adaptive systems to biologically plausible, time-dependent inputs. In an ideal situation, there would exist a one-to-one and onto map between the model parameters and the attributes that characterize the adaptive patterns (e.g., steady-state, time constants, peak values and times) so that the latter can be uniquely identified from the former. However, the presence of degeneracy [20, 21, 48], where multiple combinations of parameter values produce identical output patterns leads to the problem of unidentifiability in parameter estimation [51–55] from the available experimental or observable data.

We set out to systematically understand the structure of degeneracy in minimal models of NFBLs and IFFLs and the conditions under which degeneracy in the adaptive patterns is inherited to the impedance amplitude and phase profiles, which characterize the response of these circuits to oscillatory inputs at all frequencies, and therefore characterize the response of these circuits to a large class of time-dependent inputs, including white noise. We used analytical calculations, regular perturbation analysis, dynamical systems tools and numerical simulations.

Our results for minimal models define scenarios and hypotheses for the systematic investigation of the mechanisms of generation of adaptive patterns, resonance/phasonance and degeneracy in more complex models involving negative feedback processing, input orchestration, and combinations of these in the presence of external fluctuations and confounding factors. Our results have also implications for the development of methods for circuit and dynamical systems reconstruction based on experimental or observational data.

While our investigation was motivated by previous work originated in the field of systems biology, similar circuit structure is present in neuronal systems. Our findings for neuronal rate models describing the evolution of the instantaneous firing rates of excitatory and inhibitory networks [37–41] receiving inputs from an upstream region raise questions for more complex networks having embedded NFBLs and IFFLs as well as more complex models of spiking neurons having the same circuit structure. Our results also highlight the possibility of a variety of possible mechanisms complementary to the classical excitation/inhibition balance [56, 57]

The investigation of the mathematically tractable linear systems allowed us to identify the principles underlying the generation of degeneracy in NFBLs and IFFLs and the persistence of the degeneracy structure in the response of these adaptive systems to oscillatory inputs and therefore to time-dependent input within a large class that includes white noise. Key to these findings is the reduction of the two-dimensional system to a second order differential equation. While the two are equivalent formulations, the transformation process reduces the number of effective parameters, which we refer to as model attributes, to the minimal possible number that allow for a one-to-one and onto link between them and the (observable) attributes characterizing the adaptive patterns (time constants and steady-state). Degeneracy results from the definition of the model attributes in terms of the larger number of model parameters. The presence of degeneracy in the impedance amplitude and phase profiles and the fact that it has the same structure as in the adaptive patterns is a direct consequence of this reduction.

Because of the success of this approach, one might be tempted to believe that further reductions will aid in process. One such approach is the widely used non-dimensionalization technique [44–47]. However, we found that dimensionless NFBLs and IFFLs fail to capture the degeneracy structure of NFBLs and IFFLs both within and across circuit types. In other words, degeneracy becomes hidden by the non-dimensionalization process.

Because approach discussed above is not applicable to nonlinear systems, it is not immediately clear whether and under what conditions degeneracy is present in nonlinear models of NFBLs and IFFLs. A regular perturbation analysis of weakly nonlinear systems showed that degeneracy within circuit types, but not across circuit types is inherited to the nonlinear NFBLs and IFFLs. Numerical simulations and phase-plane analysis established that the mechanisms underlying the generation of degeneracy in nonlinear systems are qualitatively similar among them and to the ones for the linear systems. The fact that these mechanisms are qualitatively similar for different types of nonlinear models suggest the universality of the phenomena. Further research is needed to test these ideas.

The mechanisms of generation of degeneracy both within and across circuit types involve balances between the activation of the inhibitory process and the effect it exerts on the observable variable. More specifically, degeneracy within the linear NFBLs and IFFLs is created by multiple combinations of the parameter values that control the inhibition of the observable node X (*k*_2_) by the hidden node Y and its activation by X (NFBL, *k*_3_) or the input (IFFL, *ξ*). Degeneracy across circuit types is a direct consequences of these activation-inhibition balances. The persistence of degeneracy in the impedance and phase profiles for the same combination of parameter values is consistent with previous findings linking adaptive patterns with the resonance and phasonance phenomena [30–33] and precludes the ability of time-dependent inputs to resolve the degeneracy in these adaptive patterns. While the type of balances between opposing processes are expected to produce similar patterns, it is not obvious that these balances will produce the type of robust degeneracy we found. The robustness of degeneracy decreases for nonlinear systems as compared to linear ones. Specifically, while the nonlinear models show degeneracy within circuit types, they show quasi-degeneracy, a weaker form of degeneracy, across circuit types. Quasi-degenerate adaptive patterns are almost identical, but not identical. They are almost indistinguishable, but they respond differently to external perturbations, in contrast to the fully degenerate adaptive patterns. Specifically, quasi-degeneracy is not inherited by the impedance amplitude and phase profiles. This suggests that it is easier to discriminate between circuit types than to identify the precise structure of each circuit type. This is important when designing algorithms to systematically reconstruct the systems’ structure from experimental/observable data.

In previous work [48] we have identified a generic canonical model for degeneracy in the so-called *λ*-*ω* and related models. These are two-dimensional nonlinear models producing oscillations. This type of degeneracy is different from the one described here. First, in the *λ*-*ω* degeneracy is exhibited by both state variables. Second, for the *λ*-*ω* models, degeneracy is exhibited only in the steady-state regime in contrast to the NFBL and IFFL models where degeneracy is present in both the steady-state and the transient regimes. However, oscillatory models can be constituent building blocks of complex NFBL and IFFL models. Research is required to establish the dynamics of these complex networks, their degeneracy structure and the mechanisms by which the two types of degeneracy described above interact.

NFBL and IFFL have been proposed to play significant roles in the processing of temporal information and related regulatory functions in biological and neuronal systems [7–19, 37–41, 58] (and references therein). Research is needed to examine the ideas and results discussed in this paper in models describing more complex scenarios and larger networks. Additional research is needed to design computational tools to resolve degeneracy. The simple models investigated in this paper are expected to provide ground truth scenarios to text these tools.

## Acknowledgments

HGR acknowledges support from the National Science Foundation grants DMS-1608077 and IOS-2002863. ACV acknowledges support from a Grant from the Argentine Agency of Research and Technology (PICT2019-01681). HGR is a Corresponding Investigator in CONICET, Argentina and a Graduate Faculty Member in the Behavioral Neurosciences Program at Rutgers University. HGR is also Visiting Researcher/Academic in the Neurosciences Institute and the Courant Institute of Mathematical Sciences at New York University.

## A Degeneracy in neuronal firing rate NFBL and IFFL circuits

### A.1 Neuronal rate models

The rate models describing the evolution of the instantaneous firing rates of excitatory (E) and inhibitory (I) networks (Fig. 11-A) in their simplest form [37–41] are given by

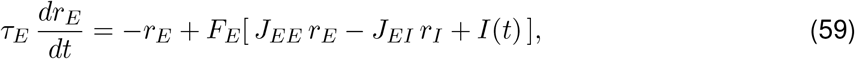

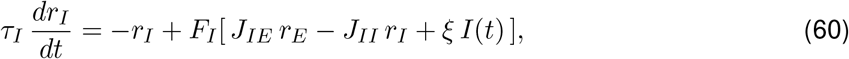

where *r*_*E*_ and *r*_*I*_ are the rates of the E- and I-nodes, *τ*_*E*_ and *τ*_*I*_ (ms) are the time constants, *I*(*t*) is the input to the E-node, *ξ I*(*t*) is the input to the I-node, and *J*_*EE*_, *J*_*EI*_, *J*_*IE*_ and *J*_*II*_ are the connectivity weights (*J*_*XZ*_ represents the connectivity weight from the Z-node to the X-node). The input-output functions *F*_*E*_ and *F*_*I*_ are monotonically increasing. Typical examples are shown in Fig. 11-B. For the NFBL rate models, *ξ* = 0, and for the IFFL rate models, *J*_*EI*_ = 0.

**Figure 11:**
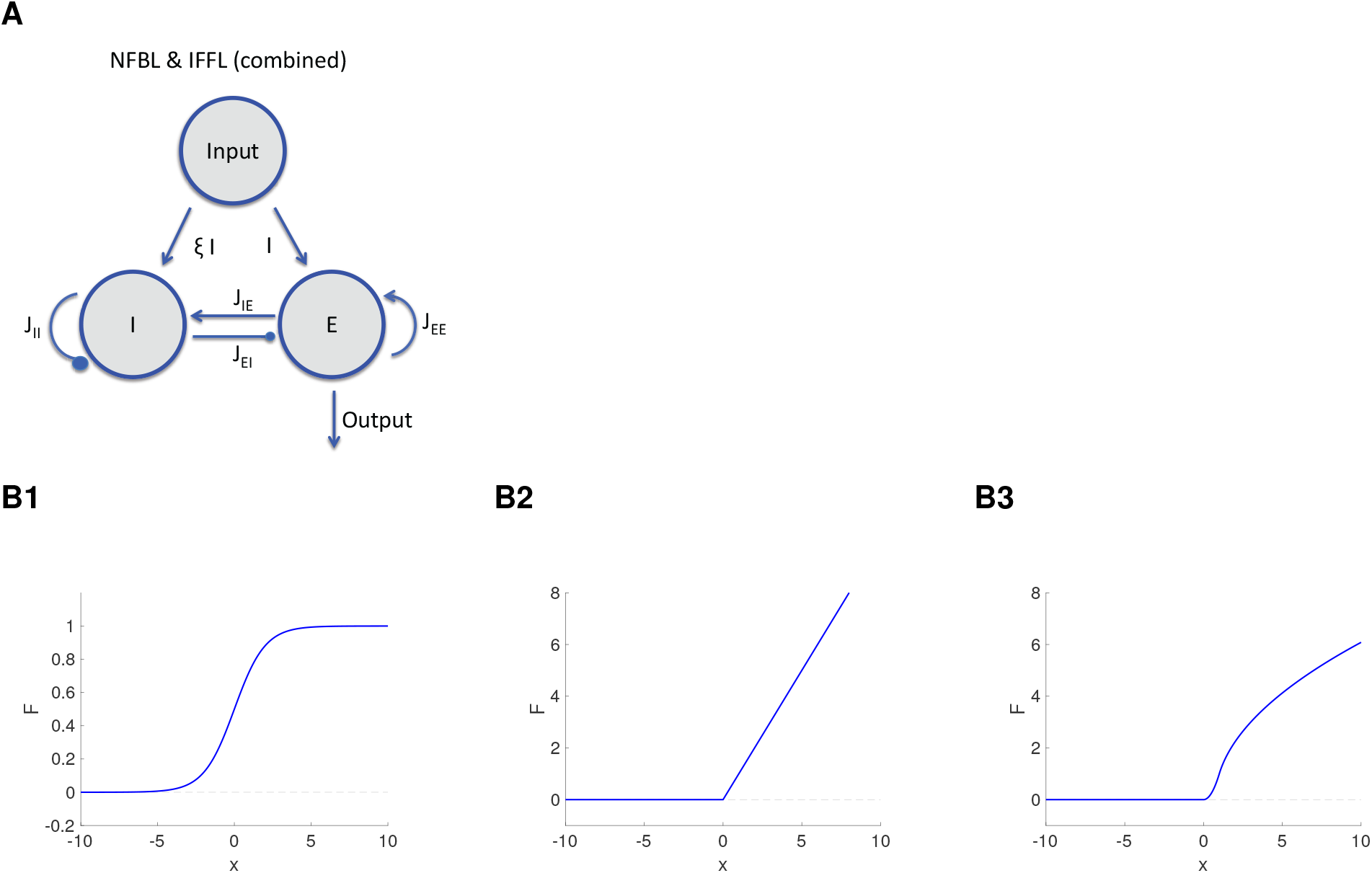
Negative feedback loop (NFBL) and incoherent feedforward loop (IFFL): schematic diagram for excitatory-inhibitory (E-I) firing rate models. **A.** Combined NFBL and IFFL circuit of a neuronal excitatory - inhibitory (E-I) network. The circuit represents a NFBL for *ξ* = 0 and a IFFL for *J*_*IE*_ = 0. The input arrives into the nodes *E* (with strength given by *I*) and *I* (with strength given by *ξ I*). The node *E* activates the node *I* (synaptic weight *J*_*IE*_), which in turn inhibits the node *E* (synaptic weight *J*_*EI*_). The two nodes also exhibit self-excitation (synaptic weight *J*_*EE*_) and self-inhibition (synaptic weight *J*_*II*_, respectively. The output is the firing rate of the node E. **B**. Representative input-output functions *F*. **B1**. Sigmoid: *F* (*x*) = (1 + *e*^−*x*^)^−1^. **B2**. Threshold-linear: *F* (*x*) = [*x*]_+_ = *H*(*x*) *x* where *H*(*x*) is the Heaviside function. **B3**. Threshold-nonlinear: 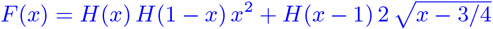.

From the topology point of view, this type of rate E-I network models are similar to the NFBL and IFFL circuits studied above. However, from the dynamics point of view, the time constants *τ*_*E*_ and *τ*_*I*_ add richness to temporal behavior of the circuits’ patterns.

### A.2 Degeneracy in the NFBL and IFFL circuits: linear neuronal rate models

From (59)-(60), the linear rate models for the excitatory-inhibitory (E-I) network (Fig. 11-A) is given by

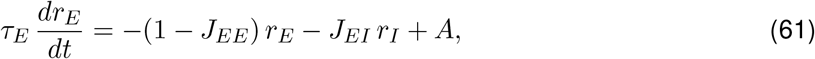

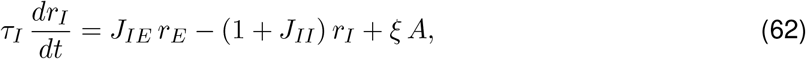

where *A* is a constant. These models can be thought of as linearization of rate models, or as describing the dynamics of threshold-linear models (Fig. 11-B2) in the region of positive values for the argument of the ReLux function.

From (11)-(12) in Section 2.2.1, system (62)-(62) can be rewritten as

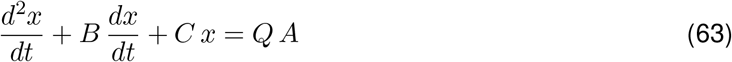

where

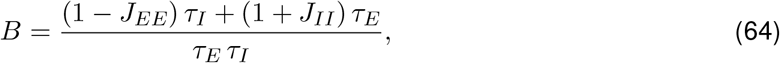

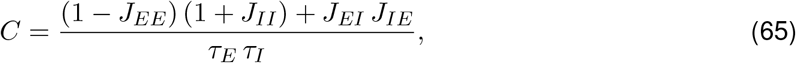

and

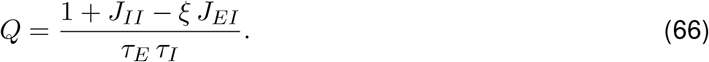

#### A.2.1 Determining the model parameters from the model attributes: NFBL

From (64)-(66) for *ξ* = 0,

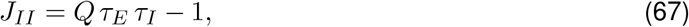

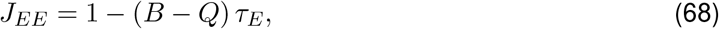

and

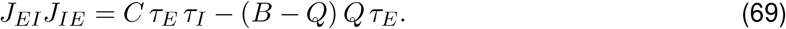

For fixed values of *τ*_*E*_ and *τ*_*I*_, *J*_*II*_ is uniquely determined by *Q* and *J*_*EE*_ is uniquely determined by *B* and *Q*. However, there are multiple combinations of *J*_*EI*_ and *J*_*IE*_ that satisfy eq. (69). This type of degeneracy is the result of the multiple balances between network excitation and inhibition as for the NFBL circuits discussed in the main body of the paper.

The presence of degeneracy within and across circuit types and its dependence on the model parameters follow from the following observations.

#### A.2.2 Determining the model parameters from the model attributes: IFFL

From (64)-(66) for *J*_*IE*_ = 0,

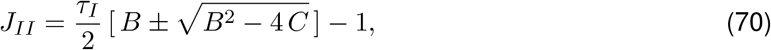

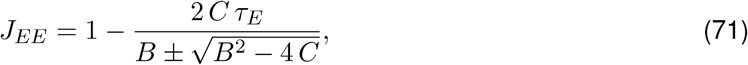

and

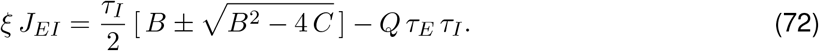

For fixed values of *τ*_*E*_ and *τ*_*I*_, *J*_*EE*_ and *J*_*II*_ are uniquely determined by *B* and *C*. However, there are multiple combinations of *ξ* and *J*_*EI*_ that satisfy eq. (72). This type of degeneracy is the result of the multiple balances between the activation of the inhibitory node and the inhibition this node exertes on the excitatory node, as for the IFFL circuits discussed in the main body of the paper.

## B. Additional analytical calculations

### B.1 Adaptive solutions to second order linear ODEs

The non-oscillatory solutions to eq. (11) for *x*(0) = 0 and *x*’(0) = *η >* 0 is given by

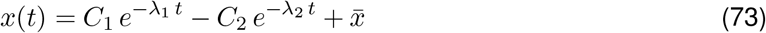

where *λ*_1,2_ = −*r*_1,2_ (17) and

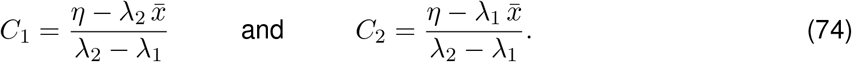

If the solution *x*(*t*) has an overshoot, then the peak time and peak value are given by

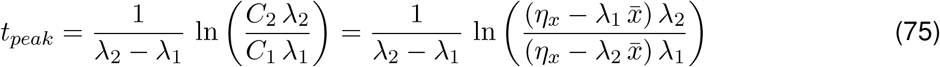

and *x*_*peak*_ = *x*(*t*_*peak*_), respectively.

The corresponding solution to eq. (13) for *y*(0) = 0 and *y*’(0) = *η >* 0 is given by

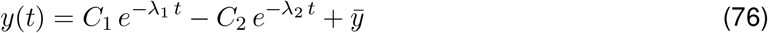

where

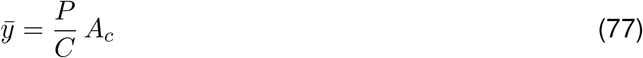

and

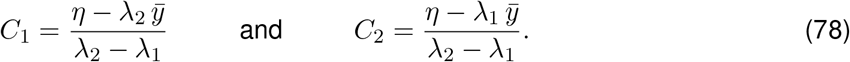

### B.2 The hidden variables corresponding to two degenerate observable variables within circuit types, but not across circuit types are multiples of each other

Degeneracy in *x* implies that *B* and *C* are fixed, but *P* is different. For two values *P*_1_ and *P*_2_ of *P*, the corresponding variables *y*_1_ and *y*_2_ must satisfy

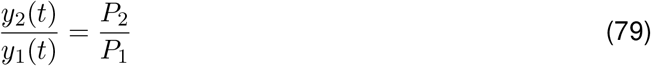

for all values of *t* provided 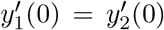 or the ratio 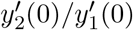 and the ratio of the steady values 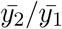 are equal (it is assumed that *y*_1_(0) = *y*_2_(0) = *y*_0_). This is the case for two degenerate NFBL adaptive patterns for which 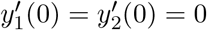 and for two degenerate IFFL adaptive patterns for which 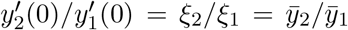 (for a IFFL, 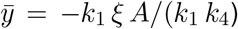). However, it is not the case across circuit types. For the NFBL, 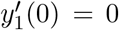, while for the IFFL,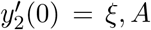. In addition, for the NFBL,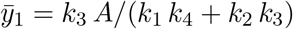, and for the IFFL, 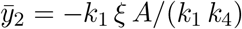.

## Supplementary Material

### S1. Recovery of the model attributes from the observable attributes

In all cases, the three model attributes (*B, C* and *Q*) obtained in Section 2.2.2 can be uniquely computed from the response pattern attributes (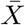, *τ*_1,2_, or, alternatively, 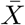, *τ* and *ω*).

Specifically for adaptive patterns, if one has access to the eigenvalues *λ*_1_ and *λ*_2_ in addition to the steady state 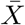, then

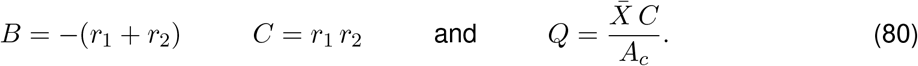

From 12, for the NFBL,

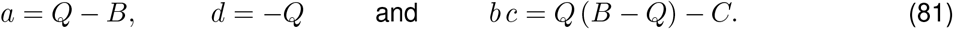

For the IFFL,

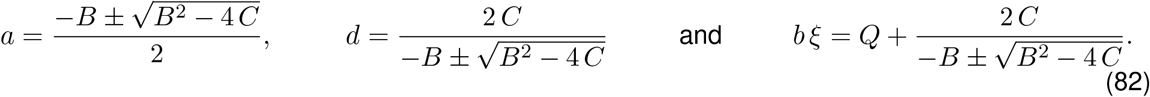

### S2. Linear systems: Impedance amplitude and phase profiles

We consider the two-dimensional linear system described by eqs. (9)-(10) in Section 2.2 with

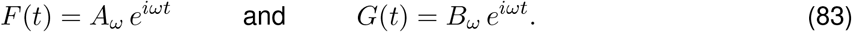

The stationary solution of system (9)-(10) is given by

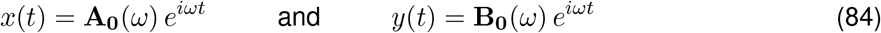

with

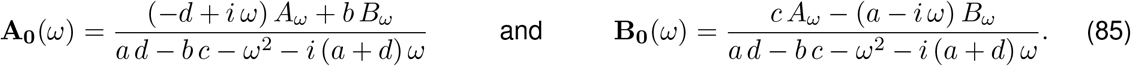

We express the input amplitudes in terms of a common parameter *A*_*in*_

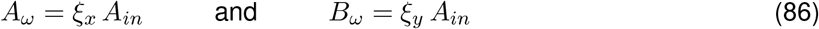

This allows us to consider inputs to only one node in the network (*ξ*_*x*_ = 0 or *ξ*_*y*_ = 0) or to both (for the NBFL-IFFL, *ξ*_*x*_ = 1 and *ξ*_*y*_ = *ξ*). By substituting into (**??**), after some algebra one obtains

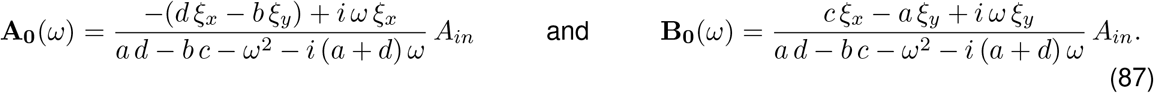

By analogy with electric circuits, we define the impedances

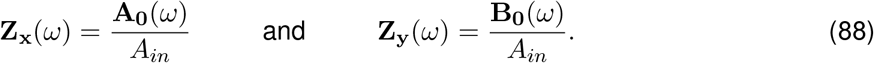

The impedance amplitudes are given by

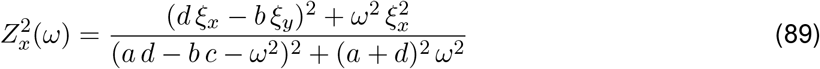

and

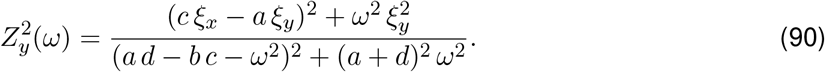

The impedance phases (phase-shift) are given by

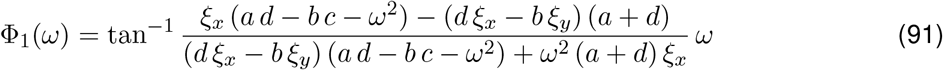

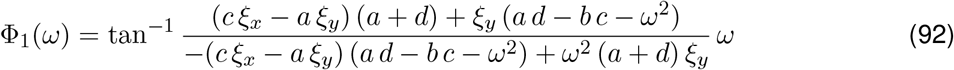

## S3. Figures

**Figure S1:**
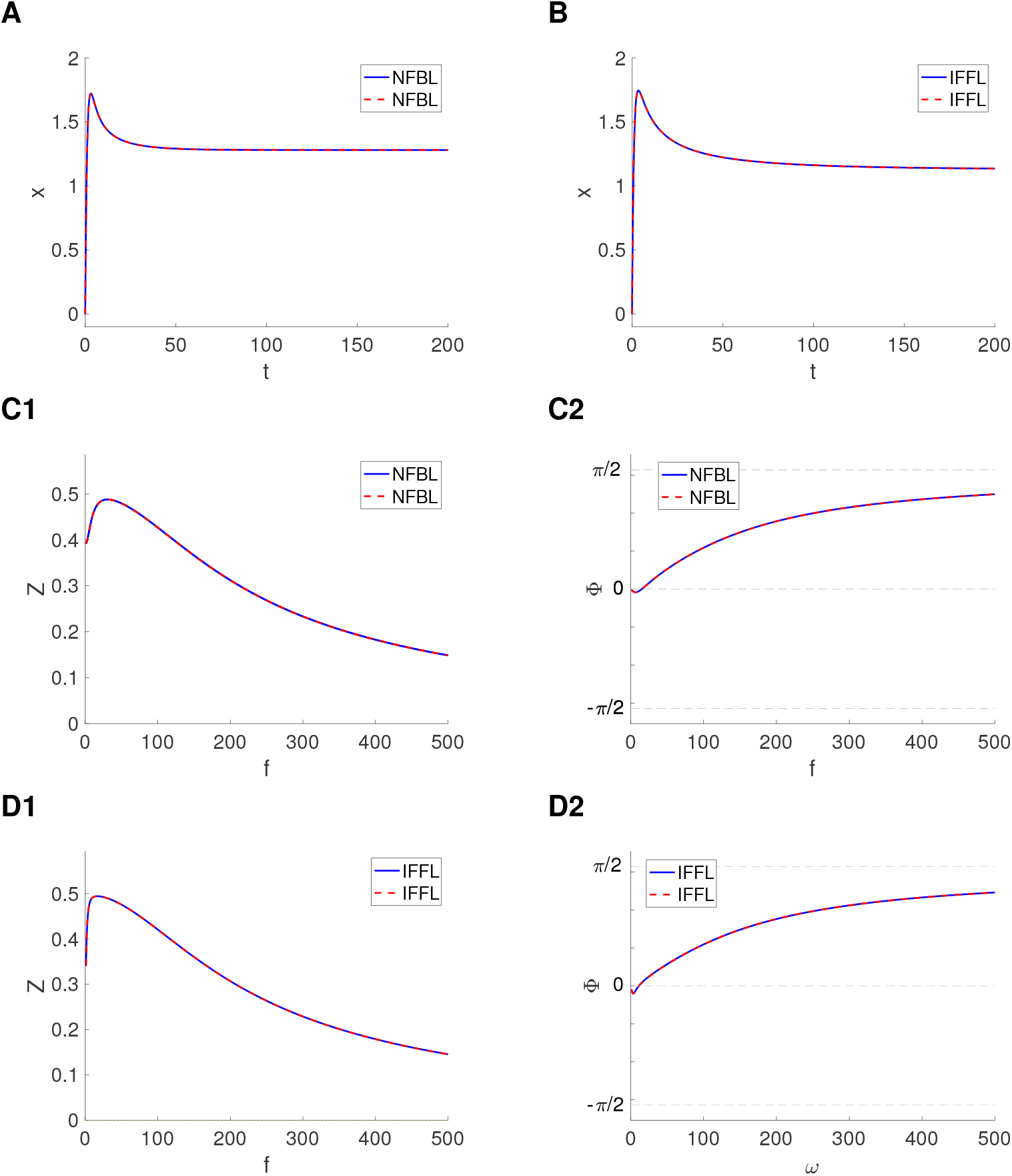
Degeneracy within circuit types in the presence of Divisive nonlinearities: representative examples or adaptive responses (to a constant input). We used System (3)-(4) with *x*(0) = *y*(0) = 0 and Γ(*x, k*_2_ *y, I*) given by eq. (7). **A, B**. Adaptive responses for *A*_*c*_ = 1 (*A*_*s*_ = 0). **C, D**. Impedance amplitude and phase profiles for *A*_*s*_ = 1 (*A*_*c*_ = 0). **A**. Parameter values: *k*_1_ = 1, *k*_2_ = 2, *k*_3_ = 0.05, *k*_4_ = 0.05, *ξ* = 0 (blue), *k*_1_ = 1, *k*_2_ = 1, *k*_3_ = 0.1, *k*_4_ = 0.05, *ξ* = 0 (dashed-red). **B**. Parameter values: *K*_1_ = 1, *K*_2_ = 2, *K*_3_ = 0, *K*_4_ = 0.015, Ξ = 0.05 (blue), *K*_1_ = 1, *K*_2_ = 0.4, *K*_3_ = 0, *K*_4_ = 0.015, Ξ = 0.25 (dashed-red). **C**. Same parameter values and color-code as in A. **D**. Same parameter values and color-code as in B.

**Figure S2:**
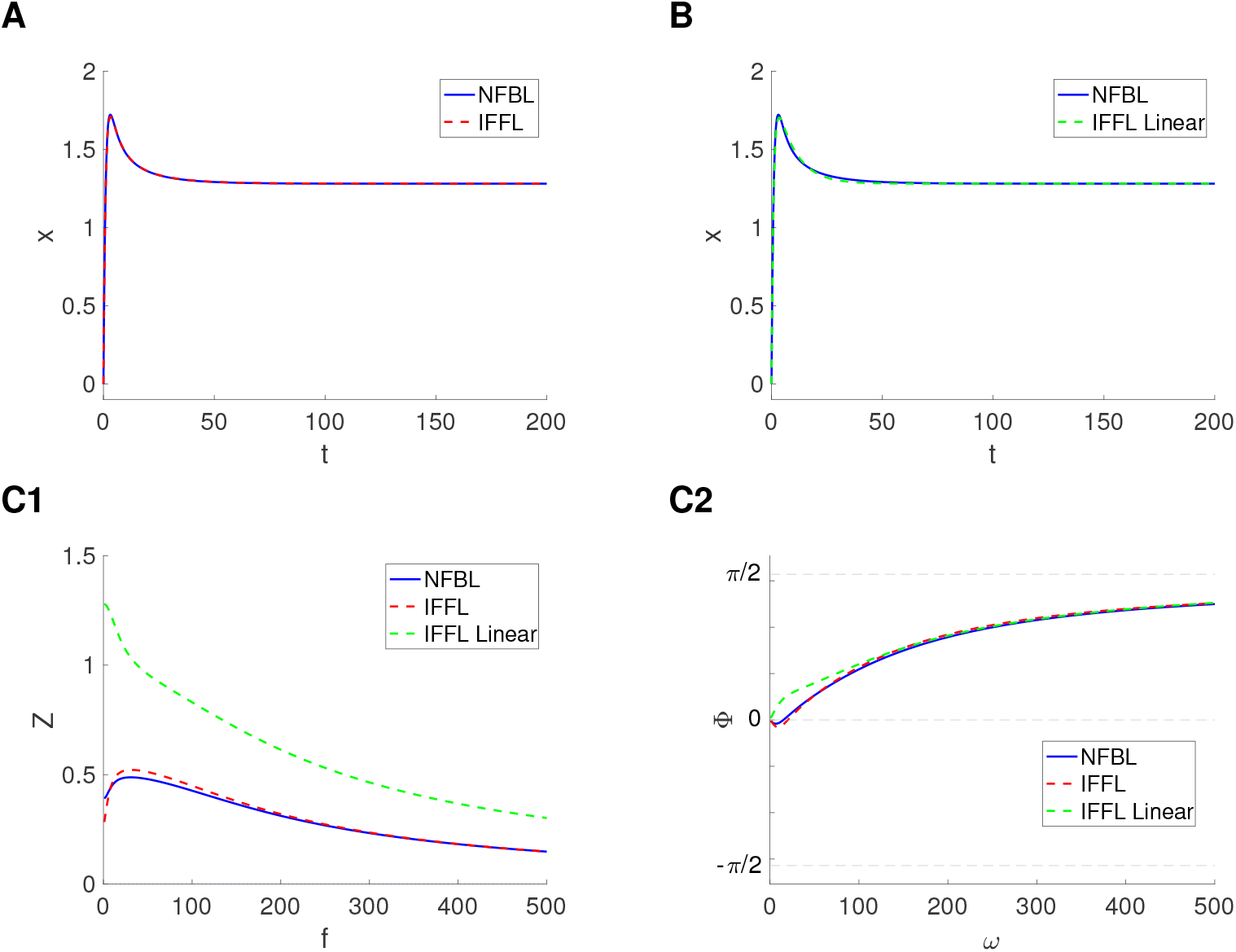
Quasi-degeneracy across circuit types in the presence of Divisive nonlinearities: representative examples of adaptive responses (to a constant input). We used System (3)-(4) with *x*(0) = *y*(0) = 0 and Γ(*x, k*_2_ *y, I*) given by (7). **A, B**. Adaptive responses for *A*_*c*_ = 1 (*A*_*s*_ = 0). **C**. Impedance amplitude and phase profiles for *A*_*s*_ = 1 (*A*_*c*_ = 0). Degeneracy is disrupted in response to oscillatory inputs. **A**. Parameter values: *k*_1_ = 1, *k*_2_ = 2, *k*_3_ = 0.05, *k*_4_ = 0.05, *ξ* = 0 (blue), *K*_1_ = 0.915, *K*_2_ = 2, *K*_3_ = 0, *K*_4_ = 0.0541, Ξ = 0.1303 (dashed-red). **B**. Parameter values: *k*_1_ = 1, *k*_2_ = 2, *k*_3_ = 0.05, *k*_4_ = 0.05, *ξ* = 0 (blue). For the linear model we used System (1)-(2) with *K*_1_ = 1.0455, *K*_2_ = 2, *K*_3_ = 0, *K*_4_ = 0.1161, Ξ = −0.0197. For these parameter values, the eigenvalues are given by *r*_1_ = −0.1161 and *r*_2_ = −1.0455. **C**. Quasi-degeneracy breaks in response to oscillatory inputs. The parameter values and color-code are as in A and B.

**Figure S3:**
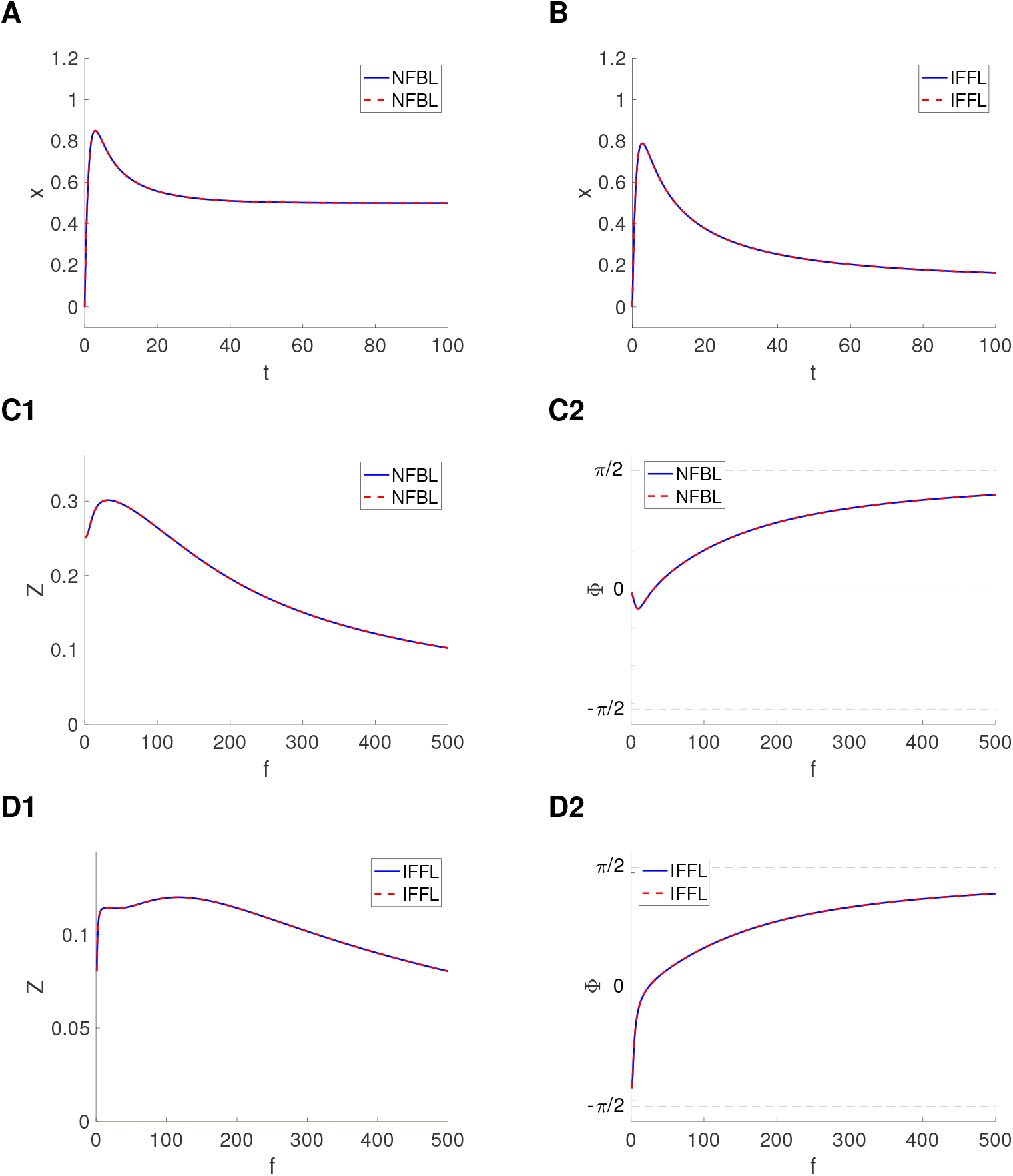
Degeneracy within circuit types in the presence of Divisive nonlinearities: representative examples or adaptive responses (to a constant input). We used System (3)-(4) with *x*(0) = *y*(0) = 0 and Γ(*x, k*_2_ *y, I*) given by eq. (8). **A, B**. Adaptive responses for *A*_*c*_ = 1 (*A*_*s*_ = 0). **C, D**. Impedance amplitude and phase profiles for *A*_*s*_ = 1 (*A*_*c*_ = 0). **A**. Parameter values: *k*_1_ = 1, *k*_2_ = 2, *k*_3_ = 0.05, *k*_4_ = 0.05, *ξ* = 0 (blue), *k*_1_ = 1, *k*_2_ = 1, *k*_3_ = 0.1, *k*_4_ = 0.05, *ξ* = 0 (dashed-red). **B**. Parameter values: *K*_1_ = 1, *K*_2_ = 2, *K*_3_ = 0, *K*_4_ = 0.015, Ξ = 0.05 (blue), *K*_1_ = 1, *K*_2_ = 0.4, *K*_3_ = 0, *K*_4_ = 0.015, Ξ = 0.25 (dashed-red). **C**. Same parameter values and color-code as in A. **D**. Same parameter values and color-code as in B.

**Figure S4:**
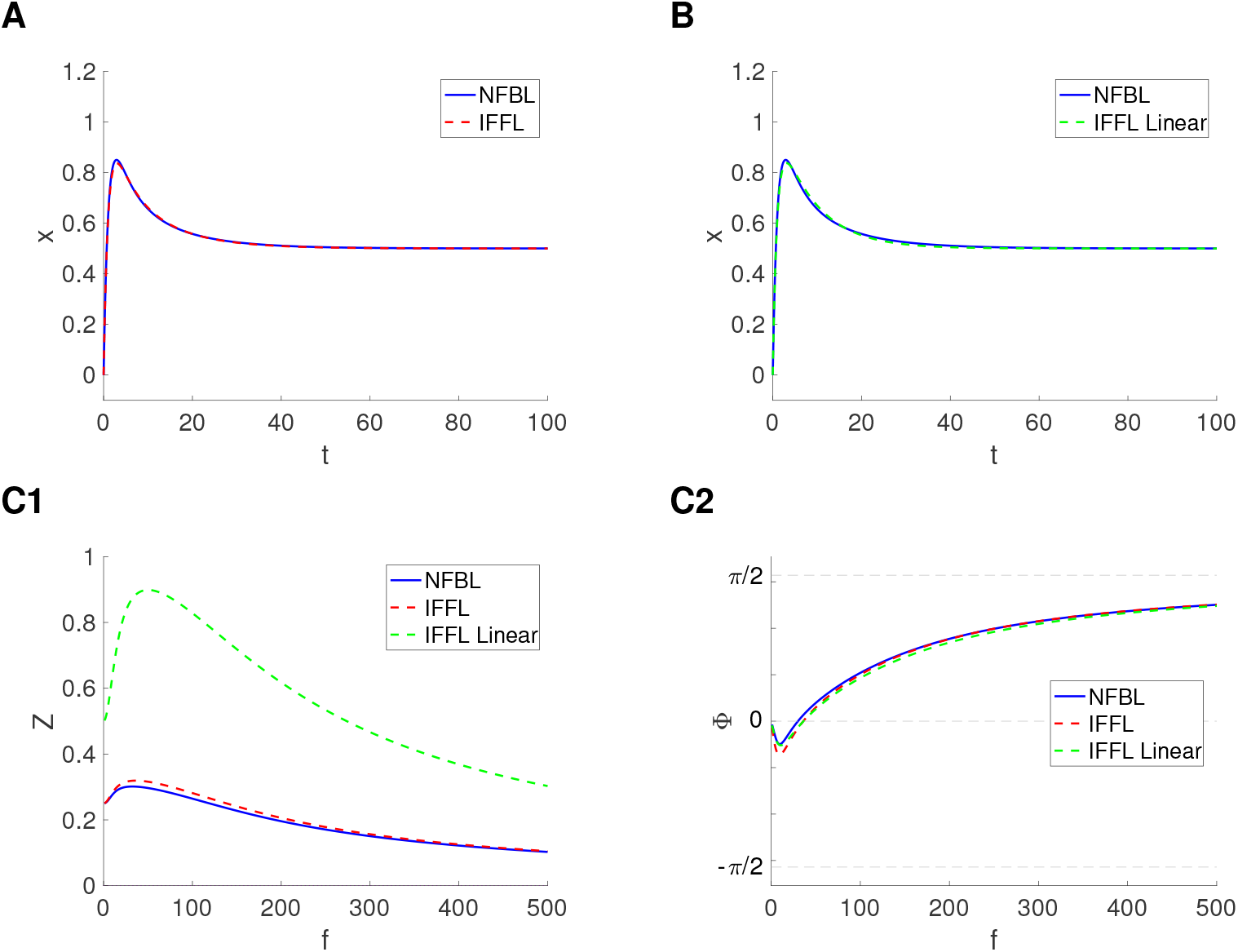
Quasi-degeneracy across circuit types in the presence of Divisive nonlinearities: representative examples of adaptive responses (to a constant input). We used System (3)-(4) with *x*(0) = *y*(0) = 0 and Γ(*x, k*_2_ *y, I*) given by (8). **A, B**. Adaptive responses for *A*_*c*_ = 1 (*A*_*s*_ = 0). **C**. Impedance amplitude and phase profiles for *A*_*s*_ = 1 (*A*_*c*_ = 0). Degeneracy is disrupted in response to oscillatory inputs. **A**. Parameter values: *k*_1_ = 1, *k*_2_ = 2, *k*_3_ = 0.05, *k*_4_ = 0.05, *ξ* = 0 (blue), *K*_1_ = 0.9594, *K*_2_ = 2, *K*_3_ = 0, *K*_4_ = 0.0855, Ξ = 0.0464 (dashed-red). **B**. Parameter values: *k*_1_ = 1, *k*_2_ = 2, *k*_3_ = 0.05, *k*_4_ = 0.05, *ξ* = 0 (blue). For the linear model we used System (1)-(2) with *K*_1_ = 1.0129, *K*_2_ = 2, *K*_3_ = 0, *K*_4_ = 0.1178, Ξ = 0.0291. For these parameter values, the eigenvalues are given by *r*_1_ = −0.1178 and *r*_2_ = −1.0129. **C**. Quasi-degeneracy breaks in response to oscillatory inputs. The parameter values and color-code are as in A and B.

## Footnotes

1 e.g.., see [48], Figs. A1-B1 and A1-B3, Appendix

2 e.g., see [48], Fig. A1-B2, Appendix

